# Tissue-adapted Tregs harness inflammatory signals to promote intestinal repair from therapy-related injury

**DOI:** 10.1101/2024.10.21.617518

**Authors:** Julius C. Fischer, Sascha Göttert, Paul Heinrich, Caroline N. Walther, Kaiji Fan, Gabriel Eisenkolb, Sophie M. Nefzger, Maximilian Giller, Omer Khalid, Vincent Ruben Timnik, Sebastian Jarosch, Lena Klostermeier, Thomas Engleitner, Nicholas Strieder, Claudia Gebhard, Sarah Diederich, Nicole Alisa Schmid, Laura Lansink Rotgerink, Laura Joachim, Sakhila Ghimire, Marianne Remke, Katja Steiger, Rupert Öllinger, Roland Rad, Daniel Wolff, Markus Feuerer, Petra Hoffmann, Matthias Edinger, Michael Rehli, Markus Tschurtschenthaler, Oliver Kepp, Guido Kroemer, Erik Thiele Orberg, Stephanie E. Combs, Wolfgang Herr, Florian Bassermann, Dirk H. Busch, Ernst Holler, Simon Heidegger, Hendrik Poeck

## Abstract

Intestinal stem cells (ISC) promote tissue repair after genotoxic or immune-mediated injury. However, ISCs are particularly sensitive to various stressors and primary targets of overwhelming immune responses such as interferon-γ (IFNγ)-mediated killing. In mouse models of gut damage and biopsies from patients having undergone allo-hematopoietic stem cell transplantation, we observed IFNy expression by intestinal T_reg_ cells. T_reg_ cells leverage combined IFNγ and interleukin 10 (IL-10) stimulation of ISCs to nurture the growth of intestinal organoids through the activation of the mTORC1 and Myc pathways. Similarly, T_reg_ cells or the combined addition of recombinant IFNγ and IL-10 promote the regeneration of organoids after irradiation. Exposure of organoids to Wnt- or EGF-free culture conditions revealed distinct growth factor-like properties of IFNγ and IL-10. While IFNγ induced epithelial proliferation and differentiation, combined addition of IFNγ and IL-10 led to balanced proliferation, ensuring ISC maintenance. Our results uncover a context-dependent role of inflammatory signaling in ISC, through which T_reg_ cells promote epithelial repair.

## Introduction

Forkhead box P3 (Foxp3)^+^ CD4^+^ regulatory T (T_reg_) cells restrain self-reactive immune responses and limit excessive inflammation ^1^. In addition to these immunosuppressive functions, T_reg_ cells have homoeostatic functions and facilitate tissue repair ^1–8^. Research into the tissue-dependent regenerative capacity of T_reg_ cells is still in its early stages, and possible biological mechanisms by which T_reg_ cells directly stimulate tissue repair appear highly context-dependent and poorly understood ^1,3,9,10^.

Nonetheless, it has been speculated that adoptive T_reg_ cell transfer might be useful for alleviating several immunopathogenic diseases including graft-versus-host disease (GVHD), which is an often fatal complication after allogeneic hematopoietic stem cell transplantation (allo-HSCT) ^11,12^. The pathogenesis of GVHD is set in motion by the effects of pre-transplant conditioning therapy resulting in collateral tissue damage and inflammation ^12^. Subsequent alloreactive priming of donor T cells, followed by their expansion and excessive effector function, ultimately leads to clinically apparent GVHD development ^13^. Several studies demonstrated that maintaining intestinal epithelial barrier function after conditioning therapy and intestinal stem cell (ISC) promoted epithelial repair restrain the development of severe GVHD ^14–17^. In general, damage to the intestinal epithelium triggers abdominal side effects associated with various cancer therapies, including chemotherapy, immunotherapy, and radiation therapy. Thus, promoting epithelial regeneration represents an important therapeutic strategy ^18–20^.

Intestinal epithelial function and the stem cell niche including ISC properties can be studied by three-dimensional multicellular organoid cultures ^21–23^. Recently, organoids have been used to elucidate the mechanisms through which different T cell subsets and their cytokines affect the renewal and differentiation of ISCs, hence contributing to epithelial regeneration after intestinal injury ^14,16,23–26^. Among other functions, T_reg_ cells and their key cytokine IL-10 have been found to help maintain the ISC niche and its stem-like function. This contrasts with other T helper cell effector cytokines, which reduce ISC renewal and rather promote ISC differentiation ^25^. In the context of tissue injury, recent studies in mice found that allogeneic T cells invade intestinal tissue and epithelial crypts early after allogeneic bone marrow transplantation (allo-BMT) and kill ISCs via epithelial IFNγ-receptor (IFNγR) activation ^27,28^. This is in line with a series of previous studies linking IFNγ to aggravated GVHD as well as epithelial tissue damage in infectious and autoinflammatory colitis ^24,29–40^. However, in allo-BMT recipients transplanted with IFNγ deficient donor T cells GVHD was found to be aggravated ^41–47^. Hence, the role of IFNγ in intestinal tissue injury is controversial, and the integration of apparently conflicting studies into a unified model has been a challenge for decades ^24,27,29–39,41–53^.

Here we describe that infiltration of IFNγ-producing T cells into the intestinal epithelium contributes both to intestinal damage and subsequent regeneration. Single-cell RNA sequencing (scRNA-seq) analyses of murine and human intestinal T_reg_ cells revealed IFNγ expression specifically after tissue adaptation upon intestinal injury. T_reg_ cells promote repair by orchestrating concomitant IFNγ and IL-10 signaling in intestinal epithelial cells, thereby skewing toxic IFNγ effects toward regenerative signals. ScRNA-seq identified that T_reg_ cells and simultaneous IFNγ/IL-10 stimulation activate mTORC1 and *Myc* signaling in ISCs, thereby promoting regeneration. Our data identify a hitherto undefined mechanism whereby tissue-adapted T_reg_ cells integrate inflammatory signals to directly foster intestinal tissue repair.

## Results

### Excessive intestinal tissue injury and regeneration are linked by IFNy and Treg cells

To investigate the relationship between damage severity and subsequent regeneration of the ISC compartment, we performed murine allo-BMTs with varying numbers of co-transplanted allogeneic T cells. Mice that experienced the most prominent acute weight loss following BMT also showed the greatest potential for weight recovery after passing the nadir (**Fig. S1A, B**) as indicated by the positive correlation between the severity of initial weight loss and the extent of subsequent clinical recovery (**Fig. 1A**). Immunohistochemical analyses and bulk transcriptomic profiling of intestinal tissue one week after allo-BMT revealed that co-transplantation of T cells dose-dependently increased intestinal T cell homing, activation, and disease activity (**Fig. S1C, D, Table S1-3**) but also induced genes related to tissue repair (e.g., type I interferon signatures, tryptophan metabolism) (**Fig. S1E, Table S1, 2**)^16,54^. Thus, the highest dose of allogeneic T cells caused the most important weight loss, followed by the strongest regenerative response.

**Figure 1:**
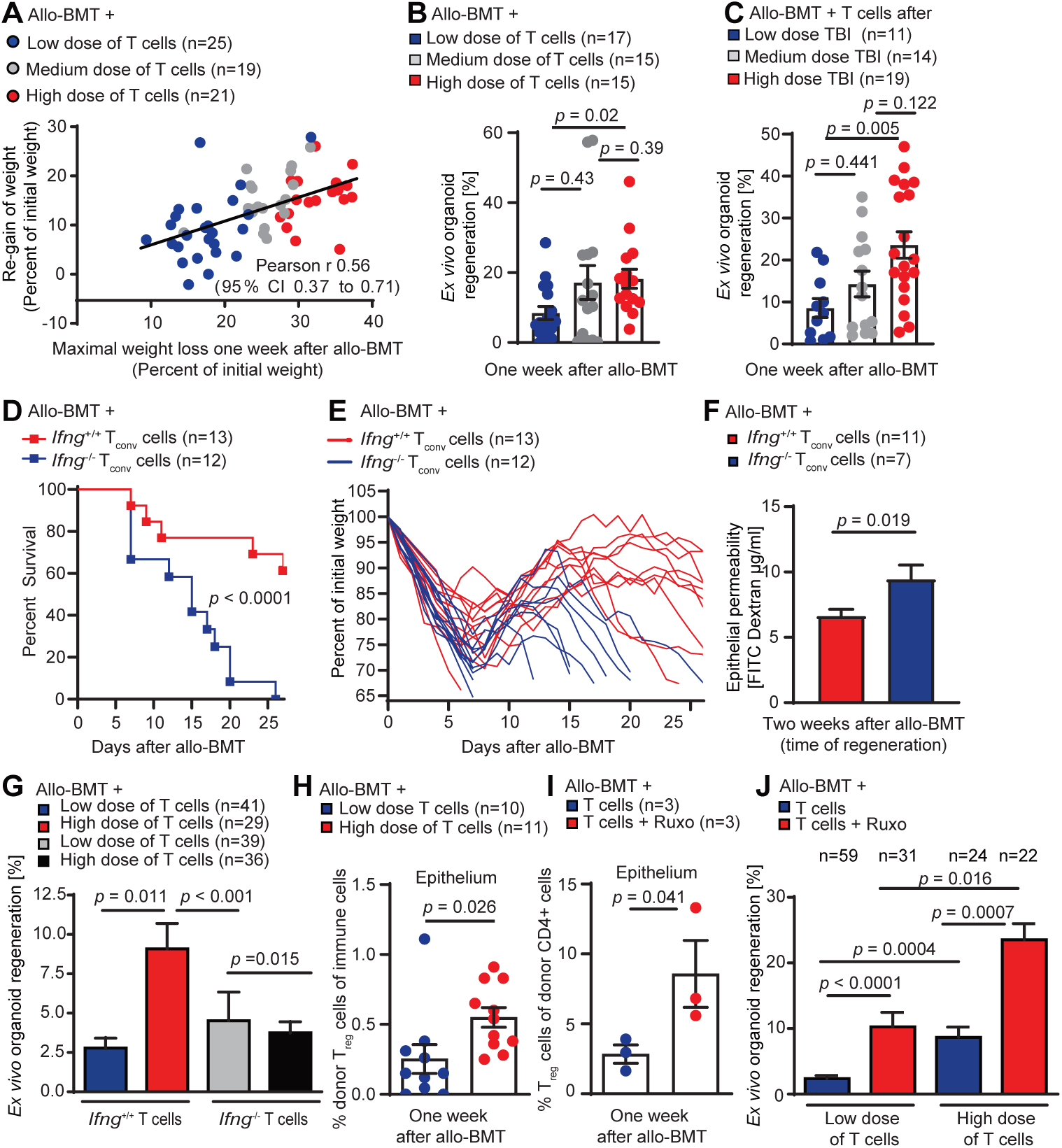
Excessive intestinal tissue injury and regeneration are linked by IFNy and T_reg_ cells. **A)** Balb/c mice received 9 Gy TBI followed by allo-BMT (C57BL/6J donors) of BM ± allogeneic T cells (low dose: 0.1 x10^6^ T cells; medium dose: 0.5 x10^6^ T cells; high dose: 2.5 x10^6^ T cells). Correlation between maximum weight loss (day 7) and peak weight recovery 2 weeks after allo-BMT. Pooled data from 3 independent experiments. **B)** Organoid regeneration rate (= number of established organoids on day 4 after starting *ex vivo* organoid culture divided by the number of cultured crypts, see Fig. S1F for details) in mice co-transplanted with increasing numbers (low, medium, high) of allogeneic T cells or **(C)** increasing doses of TBI conditioning prior to allo-BMT with medium dose of T cells. Pooled data from 3 independent experiments. **D)** Survival and **E**) weight loss of individual mice after allo-BMT with WT BM and IFNγ^+/+^ or IFNγ^-/-^ T_conv_ cells received TBI (9Gy) followed by allo-BMT (C57BL/6J donors). Pooled data from two independent experiments. **F)** Intestinal FITC dextran permeability assay two weeks (d16) after allo-BMT. Pooled data from 3 experiments. **G)** Allo-BMT with WT BM and a low or high dose of IFNγ^+/+^ or IFNγ^-/-^ T cells. *Ex vivo* organoid regeneration rate. Pooled data from 4 experiments. **H)** Intestinal intraepithelial leukocytes were isolated one week after allo-BMT and analyzed using flow cytometry after 4h of restimulation *in vitro*. Frequency of FoxP3^+^ CD4^+^ donor (H-2K^b+^) T_reg_ cells of all live CD45^+^ immune cells. **I)** Allo-BMT with BM and T cells ± ruxolitinib treatment. Depicted are intraepithelial donor T_reg_ cells. **J)** Allo-BMT with BM and a low or high dose of T cells ± ruxolitinib treatment from d-1 onwards. *Ex vivo* organoid regeneration rate. Pooled data from 4 independent experiments. Mice of the control groups with low dose of T cells and high dose of T cells are also part of Fig. 1F. Data of Fig. B, C and F-J are presented as mean ± S.E.M and were analyzed using unpaired two-tailed t-test or Kruskal-Wallis test (B, C, G, J) with Dunn’s multiple comparisons test for multiple comparisons. The number of biological replicates (n), indicating the number of mice analyzed, is shown in the figure for all individual experiments.

Given the fundamental role of the ISC niche in promoting intestinal healing, we next analyzed the capacity of the ISC compartment to support regeneration after injury (assessed by the percentage of isolated crypts that survive and form organoids during *in vitro* culture) (**Fig. S1F**). We observed that co-transplantation of higher numbers of allogeneic T cells resulted in enhanced *ex vivo* organoid regeneration (**Fig. 1B**). Similarly, increasing intestinal T cell infiltration by escalating conditioning TBI (**Fig. S1G**) induced enhanced *ex vivo* organoid regeneration (**Fig. 1C**).

Intestinal infiltration and effector function (e.g., IFNγ expression) of conventional T (T_conv_) cells is a hallmark of GVHD pathogenesis and linked to enhanced mortality^12^. We therefore performed allo-BMTs using *Ifng*^-/-^ transplants. Allo-BMT recipients co-transplanted with *Ifng*^-/-^ T_conv_ cells showed strongly reduced survival compared to mice co-transplanted with *Ifng*^+/+^ T_conv_ cells (**Fig. 1D**). Indeed, the recipients of *Ifng*^-/-^ T_conv_ cells failed to recover from acute injury during the second week after allo-BMT (**Fig. 1E**). Consistently, we observed a significant reduction in intestinal epithelial integrity (as indicated by enhanced FITC-dextran translocation from the gut lumen to the bloodstream,) two weeks after allo-BMT with co-transfer of *Ifng*^-/-^ T cells (**Fig. 1F**). Next, we studied *ex vivo* organoid regeneration of allo-BMT recipients co-transplanted with varying numbers of *Ifng*^+/+^ or *Ifng*^-/-^ T cells to explore the contribution of IFNγ to intestinal regeneration. Unlike mice co-transplanted with high doses of *Ifng*^+/+^ T cells, those receiving high doses of *Ifng*^-/-^ T cells failed to exhibit an increase in *ex vivo* organoid regeneration (**Fig. 1G**).

Collectively, these data suggest that increased intestinal T cell infiltration and IFNγ expression contribute to the capacity of the ISC niche to regenerate from excessive injury.

We next characterized the intestinal T cells infiltrate after allo-BMT and discovered that mice transplanted with high numbers of T cells showed an increased abundance of allogeneic T_reg_ cells at the peak of intestinal tissue injury (**Fig. 1H**). The Janus kinase (JAK1 and JAK2) inhibitor ruxolitinib reportedly promotes T_reg_ differentiation and ameliorates GVHD^55–57^. Accordingly, treatment with ruxolitinib increased the frequencies of epithelial T_reg_ cells following allo-BMT (**Fig. 1I**), reduced acute GVHD morbidity (**Fig. S1H**), and enhanced *ex vivo* organoid regeneration. This regenerative effect of ruxolitinib was independent of the number of co-transplanted T cells but showed the highest effect in mice that received high numbers of allogeneic T cells (**Fig. 1J**).

In summary, these results demonstrate that (i) gut-infiltrating IFNγ-expressing T cells contribute to epithelial regeneration after allo-BMT and (ii) link increased epithelial T_reg_ cell frequencies with improved regeneration of the ISC niche. By combining both aspects, we hypothesized that gut-invading and IFNγ-producing T_reg_ cells might stimulate intestinal regeneration.

### T_reg_ cells adapt to intestinal epithelial tissue injury by IFNγ expression

The adoptive transfer of donor Treg cells is a promising strategy for the prevention of GvHD. A recent study discovered that T_reg_ cells injected into mice and then recovered from different tissues (spleen, liver, large intestines) acquire organ-specific gene expression profiles within one week after allo-BMT ^58^. We re-analyzed the corresponding scRNA-seq data (**Fig. 2A**) to discover that T_reg_ cells isolated from the intestine, but not input T_reg_ cells (and less so T_reg_ cells isolated from spleen or liver) noticeably expressed IFNγ mRNA **(Fig. 2B-C**). We performed allo-BMTs with co-transplanted T cells and then employed immunofluorescence cytometry of single-cell suspensions to confirm IFNγ expression by donor T_reg_ cells in the intestinal epithelium and the lamina propria one week after allo-BMT (**Fig. S2A, B**).

**Figure 2:**
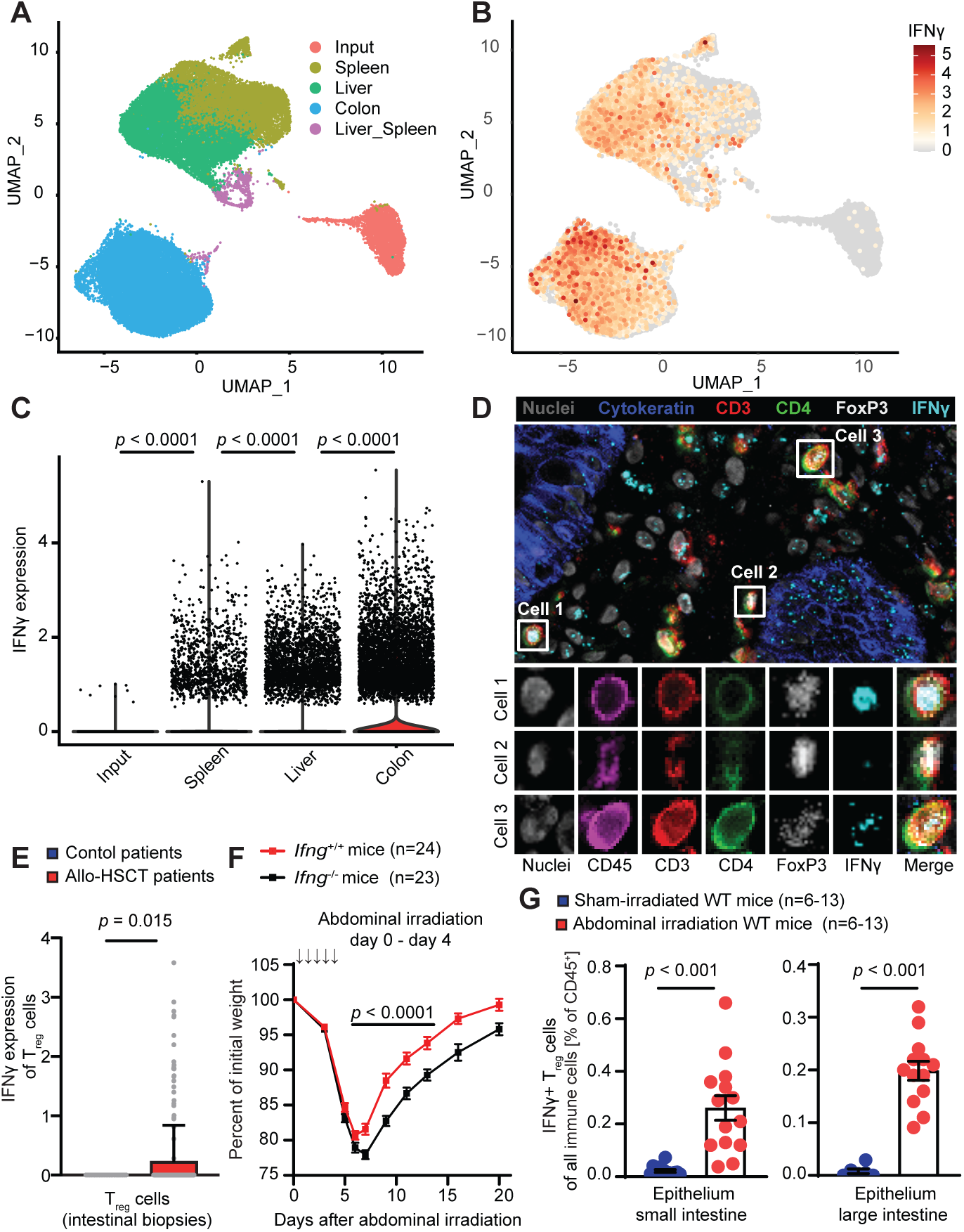
T_reg_ cells adapt to intestinal epithelial tissue injury by IFNγ expression. **A)** A published a scRNA-seq dataset (GEO accession number: GSE223798) was used^58^. Briefly, donor T_reg_ cells were expanded *in vitro*, analyzed (input T_reg_ cells) or supplied to recipient mice and then extracted from the respective tissues. Plots of single cells in UMAP space for all experimental conditions, colored by their origin or **B)** colored by their IFNγ expression. **C)** IFNγ expression of tissue and input T_reg_ cells. **D)** Exemplary image of chip cytometry of intestinal biopsies of human allo-HSCT recipients. Upper image: overlay of all indicated markers in a larger area. Lower images: high resolution of single markers and their overlay for three representative single cells. **E)** *IFNG gene* expression of T_reg_ cells was analyzed via scRNA-Seq of cells isolated from large intestinal biopsies of allo-HCST recipients (n=22 patients, with n=208 identified T_reg_ cells) or control patients that did not undergo allo-HSCT (n=5 patients with n=34 identified T_reg_ cells). **F)** WT or IFNγ^-/-^ mice received abdominal irradiation (ABI, 5x 4,5 Gy/day from day 0 until day 4) and body weight was monitored. **G)** Intraepithelial leukocytes two weeks (d15) after the start of ABI of WT mice analyzed by flow cytometry after 3h of restimulation *in vitro*. Graphs show the percentage of IFNγ^+^ T_reg_ cells of all live CD45+ immune cells. Pooled data from 3 independent experiments. Data are presented as mean ± S.E.M and were analyzed using unpaired two-tailed t-test or Mann-Whitney test (Fig. 2E), or ordinary one-way ANOVA (with Dunn’s multiple comparisons test) for multiple comparisons. The number of biological replicates (n), indicating the number of mice analyzed, is shown in the figure for all individual experiments.

We next analyzed T_reg_ cells in gastrointestinal tissue biopsies from human allo-HSCT recipients using chip cytometry ^59,60^ and identified T_reg_ cells with varying levels of IFNγ expression in close proximity to the intestinal epithelium (**Fig. 2D**). Comparative analyses of scRNA-seq data of allo-HSCT recipients and control patients corroborated the presence of intestinal T_reg_ cells with similar levels of FOXP3 expression levels (**Fig. S2C**). However, IFNγ expression was only detectable in intestinal T_reg_ cells from allo-HSCT recipients but not in T_reg_ cells from control patients (**Fig. 2E**). Thus, allo-HSCT is linked to reprogramming of intestinal T_reg_ cells towards IFNγ production.

The pathogenesis of GVHD following allo-BMT is complex as it is initiated by cytotoxic pre-transplant conditioning and exacerbated by allogeneic T cell responses. To generalize our findings, we employed a simpler model of intestinal damage relying on abdominal irradiation (ABI), which directly damages the intestinal epithelium. We found that, as compared to *Ifng^+^*^/+^ controls, ABI-treated *Ifng^-^*^/-^ mice experienced a more severe radiogenic enteritis and prolonged weight loss (**Fig. 2F**). As this occurs after allo-BMT, ABI cause T_reg_ cells to infiltrate the intestinal epithelium and to express IFNγ (**Fig. 2G, S2D-E**).

We conclude that IFNγ expression by T_reg_ cells occurs after intestinal tissue damage and then may contribute to epithelial regeneration.

### T_reg_ cell-mediated intestinal organoid growth requires epithelial IFNγ-receptor signaling

To explore the implication of IFNγ in the interaction of intestinal epithelial cells and different T cell subsets including T_reg_ cells, we cocultured intestinal organoids with T cells and then assessed the number of developing organoids after the first passage (**Fig. S3A**). Allogeneic wild-type (WT) CD4^+^CD25^-^ conventional T (T_conv_) cells impaired organoid growth. In contrast, WT CD4^+^CD25^+^ T_reg_ cells enhanced organoid growth (**Fig. 3A**). Consistent with prior studies demonstrating toxic effects of IFNγ on ISCs, IFNγ deficient (*Ifng*^-/-^) T_conv_ cells failed to reduce organoid counts (**Fig. 3B**) ^24,27^. Intriguingly, T_reg_ cell-promoted organoid growth was also IFNγ dependent, as *Ifng*^−/−^ T_reg_ cells failed to enhance organoid counts (**Fig. 3B**). Moreover, the beneficial effect of T_reg_ cells on organoid growth was abolished by neutralizing anti-IFNγ antibodies (**Fig. 3C**).

**Figure 3:**
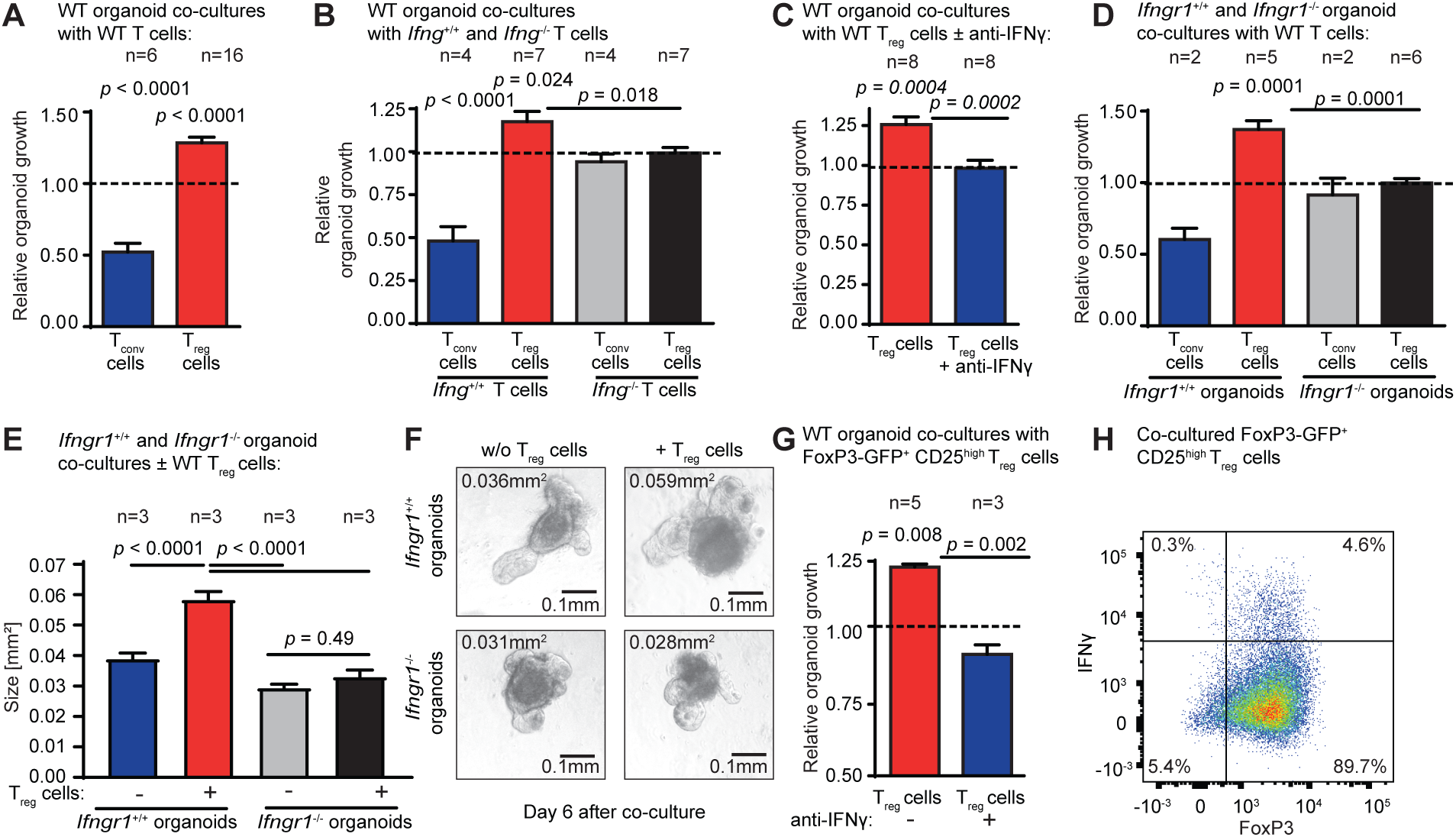
T_reg_ cell-mediated intestinal organoid growth requires epithelial IFNγ-receptor signaling. **A)** Relative organoid growth of small intestinal organoids co-cultured with allogeneic and stimulated (IL-2 + beads) CD4^+^CD25^-^ T_conv_ cells or CD4^+^CD25^+^ T_reg_ cells MACS-isolated from splenocytes of WT C57BL/6J mice (detailed description of “relative organoid growth” in methods section and Fig. S3A). The dotted line represents the growth of control organoids cultured without any T cells. **B)** Relative organoid growth of small intestinal organoids co-cultured with allogeneic T_conv_ or T_reg_ cells isolated from WT (*Ifng*^+/+^) or IFNγ-deficient (*Ifng*^-/-^) mice. Relative organoid growth of small intestinal organoids co-cultured with **C)** allogeneic WT T_reg_ cells in the presence of IFNγ blocking antibody (control organoids were cultured without any T cells but were also stimulated with IL-2 and IFNγ blocking antibody) **D)** Small intestinal organoids isolated from *Ifngr1*^+/+^ or *Ifngr1*^-/-^ mice (C57BL/6J) were co-cultured with allogeneic WT T_reg_ cells (Balb/c) to analyze relative organoid growth and **E)** mean size (area) after co-culture. Area of organoids on day 6 after co-culture; number of measured organoids: untreated *Ifngr1*^+/+^ (n=167) and *Ifngr1*^-/-^ (n=163) organoids; *Ifngr1*^+/+^ (n=116) and *Ifngr1*^-/-^ (n=113) organoids with T_reg_. **F)** Exemplary images (converted to grayscale) of organoids after co-culture. **G)** Relative organoid growth after co-culture with flow cytometry-sorted (CD4^+^ CD25^hi^ eGFP-Foxp3^+^) T_reg_ cells ± anti-IFNγ (lelt panel), and **H)** IFNγ expression of T_reg_ cells after removal from co-culture on day 4 and 4 h restimulation (eBioscience™ Cell Stimulation Cocktail plus protein transport inhibitors) (right panel). Data are presented as mean ± S.E.M and were analyzed using unpaired two-tailed t-test or ordinary one-way ANOVA (with Dunn’s multiple comparisons test) for multiple comparisons. The number of biological replicates (n), indicating the number of separate organoid culture experiments is shown in the figure for all individual experiments.

*Ifngr1*^−/−^ T_reg_ cells (which cannot respond to IFNγ due to the absence of the IFNγ receptor) retained their ability to promote organoid growth (**Fig. S3B**). In contrast, *Ifngr1*^−/−^epithelial cells failed to increase their growth in response to WT T_reg_ cells (**Fig. 3D**), meaning that the IFNγ receptor must operate in epithelial cells, not T_reg_ cells, to stimulate organoid growth. Moreover, co-culture with *Ifng*^+/+^ T_reg_ cells increased the size of developing organoids, which strictly depended on epithelial IFNγR-activation (**Fig. 3E-F**). This effect on organoid size was also lost when WT T_reg_ cells were replaced by *Ifng*^−/−^ T_reg_ cells or when IFNγ was neutralized by antibodies (**Fig. S3C**). When we increased the stringency of T_reg_ purification (by flow cytometric purification of CD4^+^CD25hi FoxP3-eGFP^+^ cells), we continued to observe IFNγ-dependent organoid growth stimulation by T_reg_ cells. This occurred in spite of the fact that only ∼5% of Tregs contained immunofluorescence-detectable IFNγ (**Fig. 3G-H**).

In summary, IFNγ plays a dual role in the intestinal response to stress. IFNγ produced by T_conv_ cells drives epithelial damage, while IFNγ produced by T_reg_ cells apparently promotes organoid growth.

### T_reg_ cell-mediated IL-10 and IFNγ co-stimulation promotes murine and human intestinal organoid growth and repair from injury

Of note, low doses of recombinant (r)IFNγ did not affect organoid growth, although it restored the ability of *Ifng*^-/-^ T_reg_ cells to induce intestinal organoid growth *in vitro*, suggesting that IFNγ is not the sole factor produced by T_reg_ cells that contributes to intestinal repair (**Fig. 4A**). We observed significantly increased epithelial permeability during the recovery phase in mice co-transplanted with Ifng^-/-^ T_conv_ cells and Ifng^+/+^ T_reg_ cells compared to recipients of both Ifng^+/+^ T_conv_ and T_reg_ cells (**Fig. 4B**). Reduced epithelial barrier function in allo-BMT recipients co-transplanted with *Ifng*^-/-^ T_conv_ cells and *Ifng*^+/+^ T_reg_ was associated with significantly reduced survival (**Fig. S4A**). We concluded that non-T_reg_ cell-derived IFNγ (T_conv_ IFNγ) is indeed important for tissue regeneration during allo-BMT. Together with our *in vitro* finding that rIFNγ was not sufficient to promote organoid growth in the absence of T_reg_ cells (**Fig 4A**), these data suggest that T_reg_ cells need to provide at least one additional signal along with epithelial IFNγR- signaling for successful intestinal recovery after allo-BMT.

**Figure 4:**
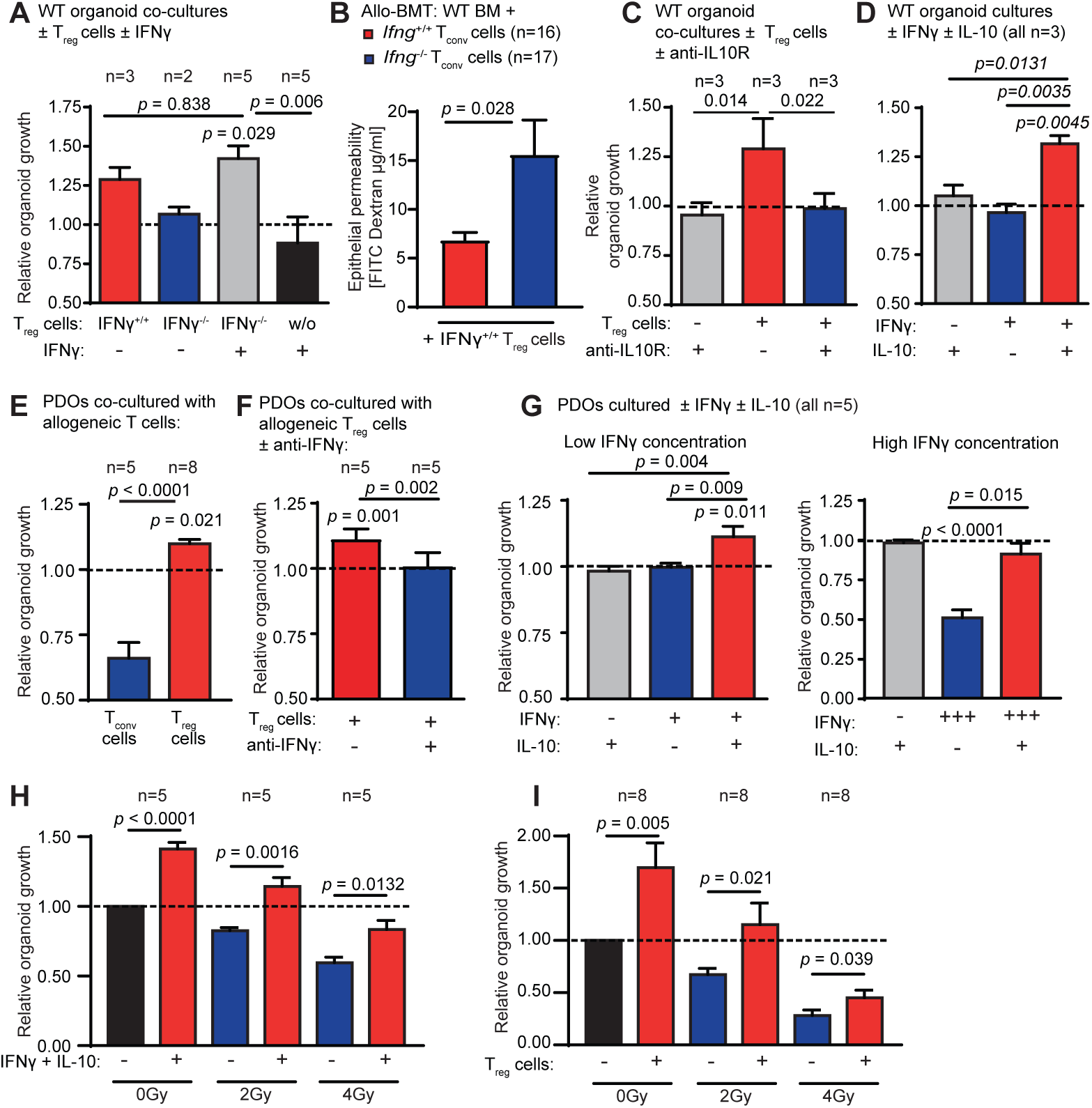
T_reg_ cell-mediated IL-10 and IFNγ co-stimulation promote murine and human intestinal organoid growth and repair from injury. **A)** Relative growth of murine small intestinal WT organoids co-cultured with allogeneic T_reg_ cells isolated from WT or IFNγ^-/-^ mice ± stimulation with 0.25 ng/mL rIFNγ. The dotted line represents growth of control organoids cultured without any T cells. **B)** Allo-BMT with WT BM plus WT T_reg_ cells and WT or *Ifng*^-/-^ T_conv_ cells. Intestinal FITC dextran permeability assay 2 weeks (d16) after allo-BMT. Pooled data from 3 experiments**. C)** Relative organoid growth of murine small intestinal WT organoids co-cultured with allogeneic WT T_reg_ cells ± anti-IL-10R antibody, or **D)** stimulated ± 10 ng/mL rIL-10 and ± 0.25 ng/mL rIFNγ in the absence of T_reg_ cells. Relative organoid growth of human patient-derived large intestinal organoids (PDOs) cocultured with **E)** allogeneic T_conv_ or T_reg_ cells, **F)** with allogeneic T_reg_ cells ± anti-IFNγ, or **G)** stimulated ± rIL-10 ± rIFNγ. Left panel: rIFNγ 0.25ng/mL; right panel: rIFNγ 5ng/mL. **H)** Relative growth of murine small intestinal organoids after *in vitro* irradiation ± cytokine stimulation, or **I)** subsequent co-culture with syngeneic T_reg_ cells. The number of separate experimental organoid cultures is indicated for all experimental conditions. All data were analyzed using two-tailed or one-tailed (Fig. 4I) unpaired t-test or ordinary one-way ANOVA for multiple comparisons (with Dunn’s multiple comparisons test) and are presented as mean ± S.E.M. The number of biological replicates (n), indicating the number of separate organoid culture experiments or the number of mice analyzed, is shown in the figure for all individual experiments.

Seeking this complementary T_reg_ cell signal that supports regeneration in IFNγR-activated organoids, we found that blocking IL-10 receptor signaling completely abolished the organoid growth-promoting effect of T_reg_ cells (**Fig. 4C).** Stimulation of intestinal organoids with the combination of rIFNγ and rIL-10 but not with either single cytokine alone was sufficient to induce organoid growth (**Fig. 4D**).

Next, we aimed at confirming our findings in a human organoid model. Therefore, we established an *in vitro* allogeneic co-culture of patient-derived intestinal organoids (PDO) and T cell subsets isolated from non-related healthy volunteers. We found that human T_conv_ cells decreased the number of co-cultured PDOs, whereas T_reg_ cells promoted organoid growth (**Fig. 4E**). In line with our murine data, T_reg_ cell-mediated organoid growth was completely dependent on IFNγ (**Fig. 4F**). Moreover, only the combined addition of rIL-10 and rIFNγ (but not that of either cytokines alone) induced human organoid growth. In addition, high rIFNγ doses resulted in direct organoid toxicity that was largely counterbalanced by co-supplemented rIL-10 (**Fig. 4G**).

Next, we characterized the role of IL-10 and IFNγ during non-T cell mediated cytotoxicity and tissue regeneration by analyzing intestinal organoids after irradiation *in vitro*. As expected, irradiation resulted in a dose-dependent reduction of organoid growth (**Fig. 4H**). Simultaneous stimulation with IFNγ and IL-10 enhanced organoid regeneration despite exposure to irradiation (**Fig. 4H**). A similar effect was observed when irradiated organoids were cocultured with syngeneic T_reg_ cells (**Fig. 4I**).

In conclusion, T_reg_ cells can promote murine and human organoid growth via combined effects of IFNγ and IL-10. Through this mechanism, T_reg_ cells stimulate recovery from immune-mediated and radiation-induced intestinal injury.

### T_reg_ cells promote organoid growth by mTORC1 and Myc activation in ISCs

To elucidate the molecular mechanisms underlying our observations, we conducted scRNA- seq of intestinal organoids i) cultured alone, ii) co-cultured with T_reg_ cells, iii) co-cultured with T_reg_ cells in the presence of IFNγ/IL-10R blocking antibodies, or iv) stimulated with rIFNγ and rIL-10.

Uniform manifold approximation and projection (UMAP) visualization yielded distinct clusters that emerged prominently after co-culturing of organoids with T_reg_ cells or or rIFNγ plus rIL-10 (Clusters 7, 8 and 12 in **Fig. S5A**). Notably, this T_reg_-induced shift was diminished when IL-10R and IFNγ signaling were inhibited. Based on a scRNA-seq reference data set from the murine intestine ^61^, we identified ISC, transit-amplifying (TA), as well as enterocyte progenitor cell subsets in clusters 7 and 12 (**Fig. 5A, S5B**). Thus, co-culture of organoids with T_reg_ cells or combined rIFNγ/rIL-10 stimulation mainly affects less differentiated cell types, which contribute to intestinal renewal and regeneration. Differentiated cell types like enterocytes, endocrine, Paneth and goblet cells were barely affected by T_reg_ cells and rIFNγ/rIL-10 (**Fig. 5A**).

**Figure 5:**
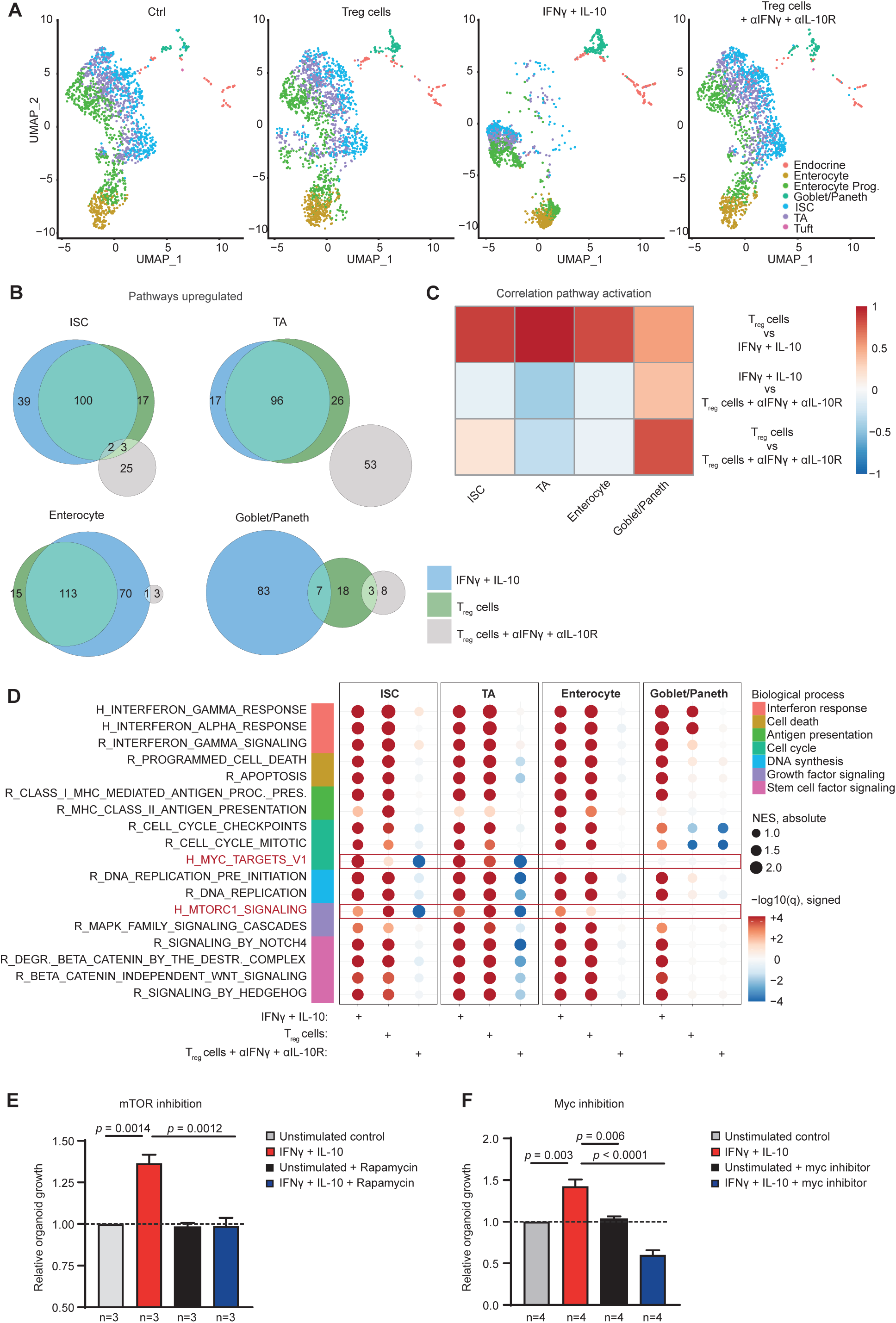
T_reg_ cells promote organoid growth by mTORC1 and Myc activation in ISCs. Murine small intestinal organoids were cocultured with CD25^high^FoxP3-GFP^+^ T_reg_ cells alone or in the presence of IFNγ and IL-10R blocking antibodies, or were stimulated with rIFNγ + rIL-10 for 4 days, and were subsequently analysed by scRNA-Seq. Pooled data from three independent experiments. **A)** Plots of single cells in UMAP space for all experimental conditions, colored by SingleR cell type annotation **B)** Venn diagrams indicating the overlap of upregulated pathways between experimental conditions compared to control organoids. Only gene sets/pathways significantly upregulated when controlling for an FDR of 10% were considered. **C)** Heatmap of Pearson correlation coefficients obtained from correlating GSEA- derived NES (normalized enrichment scores) between conditions. The values indicate correlation of regulated gene sets/ pathway activation between different conditions in indicated cell types. Only gene sets/pathways significantly up- or downregulated when controlling for an FDR of 10% were considered. **D)** Dot plot of GSEA results of selected pathways/gene sets for different cell types and treatments (vs. control). Dots are coloured according to the negative log_10_ of the GSEA q-value (FDR), the sign indicates the direction of the regulation (up positive, down negative). The size of the dots corresponds to the GSEA NES. Gene sets/pathways are derived from the Hallmark (H) and Reactome (R) gene set collections of MSigDB. **E)** Relative organoid growth of intestinal WT organoids ± stimulation with 0.25 ng/mL rIFNγ and rIL-10 and ± mTOR inhibitor rapamycin or control (DMSO) and **F)** ± myc inhibition or control (DMSO). The dotted line represents the growth of control organoids without stimulation. The number of biological replicates (n), indicating the number of separate organoid culture experiments, is shown in the figure. The data were analyzed using ordinary one-way ANOVA for multiple comparisons and are presented as mean ± S.E.M.

GSEA comparing control and various treatments across different cell types revealed that both co-culture with T_reg_ cells and stimulation with rIFNγ/rIL-10 regulated numerous distinct pathways in intestinal organoids, with a strong overlap between the two conditions (**Fig. 5B, S5C**). In line with this overlap, a high correlation of pathway regulation was observed when performing Pearson correlation analysis based on the GSEA-derived normalized enrichment scores (**Fig. 5C**). This effect was most prominent in less differentiated cell types (e.g., ISC, TA cells), perceivable in differentiated enterocytes, but undetectable in goblet/Paneth cells. In contrast, pathway regulation overlaps or correlations were scarce between, on one side, the effects of T_reg_ cell under IFNγ/IL-10 blockade and, on the other side, T_reg_ cell co-culture or rIFNγ/rIL-10 stimulation (**Fig. 5B-C**).

In search for the mechanisms of growth stimulation, we detected uniform upregulation of specific pathways in all epithelial cell types, including pathways (interferon signaling, cell death and antigen presentation) that have previously been associated with IFNγ ^27,37,62^, but also pathways associated with cell cycle regulation, DNA synthesis and growth factor signaling (**Fig. 5D, S5D**). Intriguingly, we observed a marked upregulation of mTORC1 signaling after stimulation with T_reg_ cells/cytokines, the effect induced by T_reg_ cells appearing to be more pronounced in ISC and TA cells than in enterocytes. Also, *Myc* signaling was upregulated in ISC and TA cells under these conditions, while no effect could be observed in enterocytes and goblet/Paneth cells. Both mTORC1 and *Myc* signaling upregulation was abolished when combining T_reg_ cell co-culture with rIFNγ and rIL-10 signaling blockade (**Fig. 5D**). We thus hypothesized that both pathways may play an essential role in ISC/TA cells during T_reg_ cell-mediated intestinal organoid growth stimulation. Indeed, mTOR inhibition with rapamycin or *c-myc* inhibition by compound 10058-F4 abrogated the growth-stimulating effects of IFNγ and IL-10 on intestinal organoids (**Fig. 5E, F**).

We conclude that T_reg_ cell-promoted growth of intestinal organoids is mediated by IFNγ and IL-10 through direct stimulation of ISC and cells in the early phase of differentiation (e.g. TA cells), but not through effects on mature enterocytes or other bystander cells of the ISC compartment. mTORC1 and c-*Myc* signaling are essential for these effects.

### IFNγ and IL-10 compensate for the depletion of epithelial growth factors

In the intestinal crypt, stem cell factors tightly control the proliferation and maintenance of ISCs and epithelial differentiation. Among others, epidermal growth factor (EGF) is a key regulator of proliferation and is upregulated in response to intestinal damage promoting regeneration ^63,64^. Moreover, the stem cell factor Wnt is fundamental for the maintenance and restoration of the ISC pool ^65^. We speculated that IFNγ and IL-10 might contribute to intestinal regeneration by substituting themselves to EGF and/or Wnt.

To explore whether IFNγ can compensate for the absence of EGF, we cultured murine small intestinal organoids in EGF-depleted NR media (containing Noggin and R-spondin) and simultaneously blocked endogenously produced EGF by addition of a neutralizing antibody. As expected, this resulted in reduced organoid size and numbers, compared to normal growth conditions in ENR media (containing EGF, Noggin, and R-spondin). Stimulation with IFNγ alone fully compensated for the absence of EGF. In contrast, IL-10 alone had no effect, and IL-10 also failed to augment the IFNγ effect (**Fig. 6A-C**).

**Figure 6:**
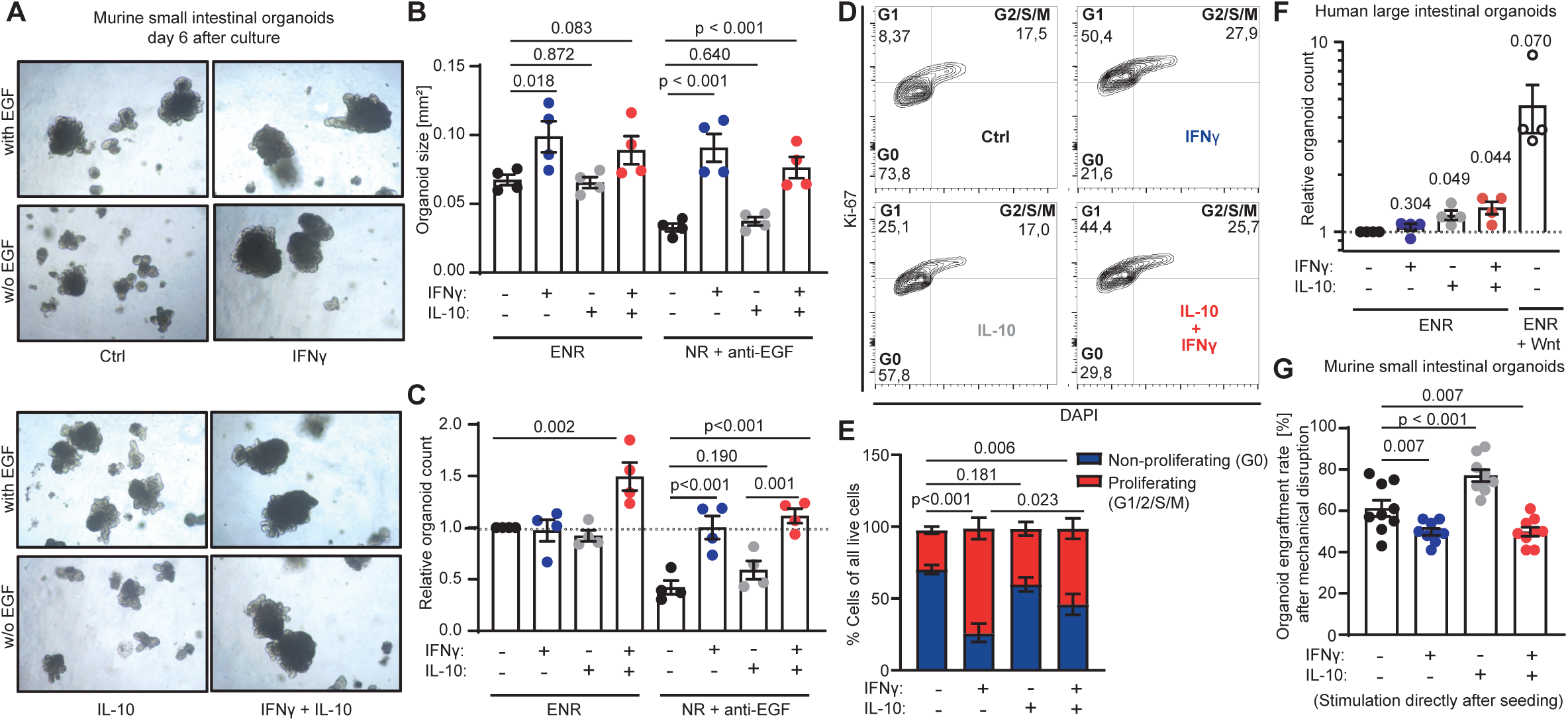
IFNγ and IL-10 compensate for the depletion of epithelial growth factors. Murine small intestinal organoids were cultured under normal growth conditions (ENR) or EGF-depleted conditions (NR + anti-EGF) and stimulated with indicated cytokines. **(A)** Representative pictures, **(B)** quantification of organoid size on day 6 of culture, and **(C)** relative organoid growth after passage (n=4 independent experiments). **(D)** Murine small intestinal organoids were stimulated for 16 h with indicated cytokines (representative plots), (**E**) the cell cycle phase was analyzed to distinguish proliferating (G1/2/S/M phase) or non-proliferating (G0 phase) cells (n=6 independent experiments). The proportion of proliferating cells was statistically analyzed. **(F)** Human large intestinal organoids were cultured under Wnt-depleted conditions and stimulated with indicated cytokines or Wnt. The number of viable organoids was determined on day 6 of culture (n=4 independent experiments). **(G)** Murine small intestinal organoids were cultured and mechanically disrupted.100 organoids were seeded into culture and immediately stimulated with indicated cytokines. The number of viable organoids was determined on day 6 of culture (n=9 wells from 3 independent experiments). All data are presented as mean ± S.E.M. p-values were calculated using ordinary one-way ANOVA plus Fisher’s LSD and one sample t-test (D).

IFNγ stimulation alone induced a rapid transition into the G1 and G2 phases compared to untreated controls (**Fig. 6D, E**). IL-10 alone did not significantly impact the cell cycle. However, when combined with IFNγ, IL-10 restricted the IFNγ-induced transition into proliferation, leading to an intermediate level of proliferation compared to controls and IFNγ alone (**Fig. 6D, E**).

Next, we sought to validate the capacity of IL-10 to support ISC retention. Since Wnt signaling is crucial for ISC maintenance in the intestinal crypt ^65^, we cultured highly Wnt-dependent large intestinal human organoids in Wnt-depleted conditions. As expected, the complete absence of Wnt drastically reduced the number of viable organoids compared to Wnt-supplemented cultures (**Fig. 6F**). This reduction was partially compensated by IL-10 stimulation, either alone or in combination with IFNγ (**Fig. 6F**).

Finally, we stimulated mechanically disrupted organoids immediately after passaging with IFNγ and/or IL-10 and determined the organoid engraftment rate (number of viable organoids grown). Stimulation with IFNγ reduced the number of engrafted organoids, while IL-10 increased this number (**Fig. 6G**). Notably, under these conditions of acute injury, the negative impact of IFNγ could not be compensated by co-administration with IL-10 (**Fig. 6G**).

In sum, these results suggest that correctly timed and dosed IFNγ can drive epithelial proliferation and organoid growth. In this context, IL-10 prevents extensive epithelial proliferation and contributes to the maintenance of the ISC pool. The combination of both properties sustainably fosters organoid growth.

## Discussion

Excessive intestinal IFNγ^+^ T cell activation is a hallmark of GVHD pathogenesis and also drives other inflammatory intestinal diseases such as immune-checkpoint inhibitor-induced colitis ^66–68^. This may be explained by the capacity of IFNγ to act kill ISC via an on-target effect on IFNγR ^24,27,28^. Here, we discovered an additional, regenerative function of IFNγ that becomes apparent in the context of simultaneous IL-10 activation.

At the cellular level, we found that T_reg_ cells provide both signals (IL-10 and IFNγ) for intestinal regeneration. While IL-10 is an established key cytokine of T_reg_ cell biology, IFNγ-producing T_reg_ cells are less well characterized ^1,3,69,70^. Here, we demonstrate that IFNγ-producing T_reg_ cells (i) promote murine and human organoid regeneration *in vitro*, (ii) invade the intestinal epithelium almost exclusively after tissue injury in mice, and (iii) locate in close proximity to the epithelial cell layer of mucosal biopsies from allo-HSCT patients, but not in healthy controls. The functional relevance of IFNγ^+^ T_reg_ cells is supported by a previous study demonstrating IFNγ expression by T_reg_ cells to be essential for the prevention of murine GVHD ^53^. Our data demonstrate that T_reg_ cells predominantly express IFNγ after adapting to intestinal tissue injury, suggesting the existence of mechanisms that cause intestinal T_reg_ cells to acquire this regenerative function. However, IFNγ produced by T_conv_ cells can contribute to intestinal repair by acting in synergy with IL-10 producing T_reg_ cells. In the same vein, a recent study reported that IFNγ produced by CD8^+^ T cells stimulates the growth of bile duct organoids ^71^.

At the level of cytokine signaling, the prominent role of IFNγ as a driver of intestinal epithelial regeneration is unexpected ^24,27^, contrasting with the established contribution of IL-10 to the prevention of inflammatory bowel disease ^72–76^. However, compared to the immunomodulatory function of IL-10 which may support intestinal repair through indirect mechanisms (such as reduced damage), its direct effects on the intestinal epithelium are poorly understood ^72–76^. Our results reveal distinct growth factor-like properties of both cytokines, which, when combined, potently promote organoid growth. Consistently, previous reports have shown that IFNγ mediates IL-10 receptor induction in epithelial cells, thus suggesting possible interactions between ^77^. While it is known that high doses of IFNγ are toxic ^24,27^, we discovered that even lower levels of IFNγ can be detrimental to the injured ISC compartment (e.g., after mechanical destruction), but strongly promote the proliferation and growth of established organoids, even compensating for EGF depletion. In contrast, IL-10 ensures protection of the ISC niche after injury but is not sufficient to induce cell cycling, which is consistent with previous reports identifying IL-10 as a key factor in maintaining the ISC pool ^25^. Building on the established knowledge that Wnt stimulation markedly improves organoid formation ^78^, we found that IL-10 compensates for Wnt depletion. Both, EGF and Wnt can be provided by Paneth cells that are often reduced upon intestinal injury, correlating with GVHD-related mortality ^16,79,80^. We conclude that the reduction of growth factor-producing bystander cells after injury can—at least in part—be compensated by combined IL-10 and IFNγ signaling. Taken together, we suggest a bivalent, dose-, timing- and context- (presence or absence of IL-10) dependent model of epithelial IFNγR signaling that can result in cell death but also favors regeneration.

On a molecular level, concurrent IFNγ/IL-10 stimulation or co-culture of T_reg_ cells with intestinal organoids promotes regenerative stimuli via mTORC1 and *Myc* signaling, particularly in ISC and undifferentiated progenitor cells. This is consistent with the established role of *Myc* in regulating epithelial cell cycle and differentiation ^81^. Consistently, we observed that IFNγ strongly promotes epithelial proliferation. The proposed pro-regenerative function of mTORC1 is in line with previous reports showing that its activation is critical for compensating damage after radiation or heat exposure ^82–84^. Along these lines, mTORC1 activation has been linked to different aspects of stem cell renewal and proliferation ^85^. However, mTOR hyperactivation in intestinal epithelial cells can also trigger necroptosis, intestinal inflammation, barrier dysfunction and cancer ^86^. At the same time, IFNγ/mTORC1 signaling controls cell death of both undifferentiated (e.g. ISC) as well as differentiated intestinal epithelial cells (e.g. Paneth cells) ^25,27,40^. Thus, the signaling strength of mTOR and/or co-activated pathways may tip the balance between tissue damage and regeneration. Our results refine this concept by demonstrating that IFNγ/mTORC1 activation stimulates ISC-mediated growth and repair in the presence of IL-10. Although we observed the most potent mTORC1 activation in ISCs and TA cells and no activation in differentiated bystander cells such as Paneth cells, we cannot formally exclude that its activation in differentiated enterocytes may contribute to enhanced epithelial regeneration. Nonetheless, it appears clear that ISC and TA cells are particularly responsive to T_reg_ cell or IFNγ/IL-10 stimulation.

Our general conclusion that T_reg_ cells promote intestinal regeneration is based on research accomplishments of the last two decades. However, T_reg_ cell-mediated tissue protection is often explained by their tolerance-inducing and immunosuppressive functions, which restrict inflammation and thus create a favorable environment for regeneration ^1^. Yet, direct links between T_reg_ cells and tissue repair remain scarce ^9,87,88^. Only a few landmark studies identified T_reg_ cells to be able to directly induce regeneration via the release of amphiregulin or jagged 1 ^2,7,8^. In the context of intestinal damage induced by allo-HSCT, T_reg_ cells were shown to restrain donor T_conv_ cell activation and proliferation in lymphoid organs, and thus reduce damage to the gut ^1,68^. However, beyond this effect, the transfer of T_reg_ cells results in enhanced signs of intestinal regeneration after allo-BMT ^89^. Additionally, T_reg_ cells can increase stemness in ISCs, help maintain the ISC niche in *vivo,* and foster intestinal regeneration via IL-10 activation during helminth and bacterial infections ^25^.

Our findings have translational implications. Using ruxolitinib to treat GVHD in mice, we were able to increase the proportion of intestinal epithelial T_reg_ cells, which was accompanied by enhanced regeneration of the ISC compartment. This is in line with our previous report of improved T_reg_ cell differentiation in ruxolitinib-treated mice after allo-BMT ^57^. Ruxolitinib is a JAK1/2 inhibitor that inhibits IFNγR/IL-10R signaling, which may appear incompatible with cytokine-mediated epithelial regeneration. However, therapeutic JAK inhibition *in vivo* most likely does not completely block epithelial cytokine signaling. Consistent with this, ruxolitinib showed the highest potential to foster intestinal regeneration in mice transplanted with high numbers of allogeneic T cells and presumably potent residual IFNγ activation. We hypothesize that enhanced ruxolitinib-promoted epithelial T_reg_ cell infiltration pave the way for optimal regenerative responses.

The implications of our findings extend beyond T cell-mediated diseases, as (i) IFNγ-deficient mice showed delayed intestinal regeneration after abdominal irradiation and (ii) T_reg_ cells and IFNγ/IL-10 stimulation promoted growth of intestinal organoids after irradiation as well. Notably, a previous report found that treatment of mice with recombinant IFNγ prior to total body irradiation resulted in aggravated intestinal injury ^24^. Taken together, these findings i) broaden the proposed regenerative mechanisms of IFNγ/IL-10 to non-immune cell-mediated forms of tissue damage, and ii) highlight context-dependent functions based on timing, signaling strength and concurrent factors.

Our study has several limitations. Thus, we provide several lines of evidence that IFNγ and IL-10 are critical for promoting intestinal regeneration and that T_reg_ cells are able to provide both signals. However, to which extent other cellular sources of IFNγ and/or IL-10 contribute to intestinal regeneration by stimulating the same epithelial pathways currently remains unclear. While we provide data that T_conv_ cell-derived IFNγ is critical for restoring epithelial barrier function *in vivo*, we can only speculate about possible non-T_reg_ cell-derived sources of IL-10 (e.g. B cells, type 2 T helper cells). Importantly, we found that rIFNγ/rIL-10 stimulation fostered epithelial regeneration, which was completely independent from any cellular source. We therefore suggest context-dependent (e.g., disease-specific) cellular sources of IFNγ/IL-10 *in vivo,* and do not postulate that only T_reg_ cells would be capable of exploiting this mechanism to stimulate intestinal regeneration.

In conclusion, our study provides a new perspective on how the ISC niche responds to immune or non-immune cell-mediated injuries and integrates different cytokine signals to direct the fate of the intestinal epithelium into tissue destruction or regeneration. While high-dose IFNγ is invariably toxic, low-dose IFNγ combined with IL-10 stimulation has a marked repair-stimulatory effect. Tissue infiltrating Treg cells can simultaneously provide both cytokines to achieve this favorable effect.

## Methods

### Human studies

Human studies were approved by the local authorities (Ethics Commission of the Technical University of Munich, School of Medicine, study number 458/17 S and the Ethics Commission of the University of Regensburg 14-101-0047). These studies included male and female patients who were included regardless of their gender.

### Mice

Animals were housed in specific pathogen-free (SPF) animal research facilities and were monitored for pathogens according to FELASA recommendations. Animal studies were approved by the local regulatory authority (Regierung von Oberbayern, Munich). C57BL/6J (H- 2kb) and Balb/c (H-2kd) mice were purchased from Janvier Labs (France). Male or female mice were between 6 and 12 weeks of age at the onset of experiments. IFNγ-deficient mice (*Ifng^-/-^)* (C57BL/6J) and IFNγ receptor-deficient mice (*Ifngr1^-/-^)* have been described previously ^90,91^. FoxP3-EGFP mice were kindly provided by Bernard Malissen and have been described previously ^92^. Transgenic mice were co-housed with age- and sex-matched WT controls after weaning for a period of 4–6 weeks before starting *in vivo* experiments.

### Induction of GVHD after allo-BMT and treatment with ruxolitinib

Induction of GVHD after allo-BMT with myeloablative TBI using major mismatch (H-2kd/H-2kb) GVHD mouse models was performed as previously described ^93^. Treatment with ruxolitinib was performed as previously described ^57^. Briefly, Balb/c recipients were intravenously injected with 5×10^6^ allogeneic (C57BL/6J donor mice) T cell-depleted BM cells directly after myeloablative TBI with 2 x 4.5 Gy (medium dose). In some experiment, mice received 2 x 4.0 Gy (low dose) or 2 x 5.0-5.5 Gy (high dose). Co-transplantation of allogeneic T cells was typically done with 0.5 x 10^6^ purified allogeneic donor T cells (medium dose). In some experiment mice received 0.1 (low dose), 1.5 - 2.5 x 10^6^ T cells respectively (high dose), as indicated in the figure legends. Details are described in the supplementary information.

### Crypt isolation

Isolation of intestinal epithelial crypts was performed as previously described ^22^. Briefly, small intestines were cut longitudinally, washed and were incubated in 10 mM ethylenediamine-tetraacetic acid (EDTA) for 25 min (4°C) to dissociate the crypts. The supernatant containing crypts was collected.

### Murine organoid culture (generation of organoids for further experiment)

Crypts were suspended in liquefied growth factor-reduced Matrigel (Corning) (33% ENR- medium or PBS; 66% growth factor-reduced Matrigel) at 4°C, and were plated in delta-surface Nunc 24-well plates in 30 μL drops, each containing approximately 200 crypts. After the Matrigel drops polymerized, 500 µL complete crypt culture medium was added to small intestine crypt cultures. ENR-medium was prepared as follows: advanced DMEM/F12 (Life technologies), 2 mM L-glutamine (Sigma), 10 mM HEPES (Life technologies), 100 U/m penicillin, 100 μg/mL streptomycin (Life technologies), 1.25 mM N-acetyl cysteine (Sigma), 1x B27 supplement (Life technologies), 1x N2 supplement (Life technologies), 50 ng/mL mEGF (Peprotech), 100 ng/mL rec. mNoggin (Peprotech), 10% human R-spondin-1-conditioned medium of hR spondin-1 transfected HEK 293T cells. All plates were incubated at 37 °C / 5% CO_2_ and medium was replaced every 2-3 days. After 7 days organoids were passaged by mechanical disruption with a 5 mL syringe and an 18 G needle.

### *Ex vivo* murine organoid culture (assessment of *ex vivo* organoid regeneration)

The isolation of murine small intestinal crypts was carried out as described above (section “Murine Organoid Culture”). Crypts were isolated on day 7 after allo-BMT during acute intestinal tissue injury, counted, and organoid culture was started with a defined number of crypts (n = 200/drop). At least two drops per mouse were generated (=technical replicates). The number of viable organoids was counted on day 3 after seeding, divided by the number of crypts used, and reported in the figures as a percentage of initial crypts (x / 200 x 100%). Fig. S1F illustrates the experimental setup.

### Human organoid culture

For human organoids, healthy tissue of colon resections of colorectal cancer patients was used. The experiments were performed in a similar manner to the murine experiments but using specific human culture conditions, which are described in detail in the supporting information.

### Co-culture of organoids and allogeneic T cells

PBMCs from healthy human donors were isolated with Biocoll cell separation solution. T_reg_ cells were then isolated from PBMCs using the CD4^+^CD25^+^ Regulatory T Cell Isolation Kit (Miltenyi Biotec) according to the manufacturer’s protocol. Murine T cells were isolated from C57Bl6/J (WT or IFNγ^-/-^) splenocytes with CD4^+^CD25^+^ Regulatory T Cell Isolation Kit (Miltenyi Biotec) according to the manufacturer’s protocol. CD4^+^CD25^+^ cells were used as T_reg_ cells, CD4^+^CD25^-^ cells as conventional T cells. For coculture, species-matched allogeneic T cells were added to passaged human organoids or passaged Balb/c small intestinal organoids with either human IL-2 (30 U/mL, Peprotec) and Dynabeads human T-activator CD3/CD28 (ThermoFisher, 2 µL per well), or murine IL-2 (30 U/mL, Peprotec) and Dynabeads murine T-activator CD3/CD28 (ThermoFisher, 2 µL per well) respectively. To prevent direct contact between organoids and T cells, plates were kept slightly tilted with the organoid drop above the T cells. T cells were added in concentrations as indicated in the figure legends. Blocking antibodies (InVivoMAb anti-mouse IL-10R (CD210), InVivoMAb anti-mouse IFNγ, InVivoMAb anti-human IFNγ, all Bioxcell) were added in a concentration of 10µg/mL at onset of the coculture. Recombinant IFNγ (Recombinant Murine IFN-γ, Peprotec) was added at onset of indicated cocultures with IFNγ^-/-^ T_reg_ cells in a concentration of 0.25ng/mL. After 4 days, T cells were removed. Photos for size determination were taken at day 6 and area size of organoids was analyzed in a blinded fashion using the ImageJ-based software Fiji. Organoids were passaged after 7 days and counted on day 3 after passage.

### Stimulation of intestinal organoids

For cytokine stimulations, human or murine organoids were seeded. After 24 h, 0.25 ng/mL or 5 ng/mL recombinant IFNγ (recombinant murine Ifnγ or recombinant human IFNγ, both Peprotec), 10 ng/mL recombinant IL-10 (recombinant human IL-10 or recombinant murine Il-10, both Peprotec) or a combination of both were added to the culture. The mTOR inhibitor Rapamycin (1 µg/mL, Invivogen) or c-myc inhibitor 10058-F4 (100 µg/mL, Sigma-Aldrich) was added 6 h after organoid seeding. Cytokines and inhibitors were removed after 4 days. Photos for size determination were taken at day 6 and the area of organoids was analyzed in a blinded fashion using the ImageJ-based software Fiji. Organoids were passaged after 7 days and counted on day 3 after passage.

### Organoid engraftment experiments

Organoids were cultured and passaged as described above (murine organoid culture). After mechanic disruption of grown organoids, newly seeded cells were directly stimulated with 0.25 ng/mL rIfnγ and/or 10 ng/mL rIl-10.

### Abdominal irradiation

Specific anatomic regions of mice were irradiated as previously described ^94^. Details are described in the supplementary information.

### Irradiation of intestinal organoids and co-culture with T_reg_ cells or cytokine stimulation

Intestinal organoids were cultured and passaged as described above and irradiated using previously described techniques ^95^.

### Statistical analysis

Animal numbers per group (n) and/or number of co-culture experiments (n) are depicted in the figures. GraphPad Prism was used for statistical analysis. Applied statistical tests for all individual experiments are indicated in the figure legends. Survival was analyzed using the Log-rank test. Normal distribution was assumed and differences between means of two experimental groups were analyzed using unpaired t-test. Two-sided tests were always used unless the experiments were based on previous ones with significant results after two-sided testing (Fig. 4I). We did not assume a normal distribution in ex vivo organoid cultures and ex vivo scRNA analyses, which clearly showed non-normally distributed patterns and Mann– Whitney U test and Kruskal-Wallis test was used. We used ordinary one-way ANOVA and Kruskal-Wallis test for multiple comparisons and performed Dunnett’s test for Multiple-test corrections. Multiple testing corrections were omitted in individual experiments that consisted of multiple previously investigated experimental conditions. P values are shown in the figures. Further details, including statistical and methodological details of scRNA-Seq and bulk tissue RNA-Seq analyses, are provided in the supplementary materials.

## Supplemental information

I. Additional Material and Methods
II. Table S4 (Antibodies)
III. Supplemental Figures 1-6:

1. Intestinal T cell infiltration determines acute weight loss and regeneration after allo-BMT
2. Enhanced intestinal abundance of IFNγ-expressing T_reg_ cells after tissue injury
3. T_reg_ cell-derived IFNγ promotes the growth of intestinal organoids independently of an IFNγR feedback loop on T_reg_ cells
4. T_conv_ cell-derived IFNγ is essential for T_reg_ cell-mediated protection from GVHD after allo-BMT
5. Clustering, cell type annotation and overlap of downregulated signaling pathways between experimental conditions
6. UMAP plots of UCell scores and pots of single cells in UMAP space of single cell RNA sequencing analysis of murine small intestinal organoids
IV. References of supplemental information

## Acknowledgement

We thank Tatiana Nedelko and Maria Krieger (Poeck group and Heidegger group) as well as Olga Seelbach and Marion Mielke (Comparative Experimental Pathology, School of Medicine, TUM) for excellent technical support. We thank Hannah Felchle (Fischer group) for her help monitoring animal experiments. Sequencing was performed at TUM (Roland Rad group) and the NGS Core of the LIT (Regensburg, Germany). We thank Hanna Stanewsky for technical support.

## Author contributions

J.C.F. initiated the study, designed, established, conducted and analyzed experiments, wrote the manuscript and acquired funding. S.G. designed, established, conducted and analyzed experiments and wrote the manuscript. G.E. designed, conducted and analyzed initial experiments. C.N.W., P.H., K.F., S.M.N., M.G., V.R.T., O.K. S.J., T.E., R.Ö., E.T.O., M.R. and K.S. performed and analyzed experiments. S.D., L.L.R., L.K., L.J., N.A.S., N.S., C.G. and S.G. helped to perform and analyze experiments. R.R., P.H., D.W., M.F., M. E., M. R., M.T., O.K., S.E.C., W.H., F.B. provided intellectual input. E.H. and D.B. guided the analysis of the human GVHD samples. H.P., G.K. and S.H. analyzed the data, edited the manuscript, and acquired the funding. H.P. and J.C.F. directed the study. This work is part of the doctoral thesis of S.G. at UKR/TUM and C.N.W. at TUM.

## Data and materials availability

### Lead contact

All requests will be fulfilled by the lead contact Hendrik Poeck (E-mail address).

### Materials availability

This study did not generate new unique reagents.

### Data availability

All sequencing data is publicly available at the time of publication. Single cell RNA-Seq data of murine organoids were deposited at Gene Expression Omnibus (GEO) (accession number: GSE252335). The scripts used for scRNA-Seq analysis have been deposited to Github (https://github.com/lit-regensburg/scRNA_intest_org_co-culture/). RNA- Seq. data of bulk murine tissue were deposited at European Nucleotide Archive (ENA) (accession number PRJEB71507). Processed RNA-sequencing data and lists of differentially expressed genes, are available in the Supplementary Materials (Table S1-S3). This publication includes additional analysis of scRNA-Seq data of human samples that were previously described ^60^ and were deposited at (GEO) (accession number: GSE234357) and previously described^58^ murine data (GEO accession number: GSE223798). Any additional information is available from the corresponding authors upon reasonable request.

