## Supplemental information for "Tissue-adapted Tregs harness inflammatory signals to promote intestinal repair from therapy-related injury"

**Contents of this file:**

I. Additional Material and Methods
II. Table S4 (Antibodies)
III. Supplemental Figures S1-S6
IV. References of supplemental information

#### I. Additional Material and Methods

##### Induction of GVHD after allo-BMT and treatment with ruxolitinib

Induction of GVHD after allo-BMT with myeloablative TBI using major mismatch (H-2kd/H-2kb) GVHD mouse models was performed as previously described [1]. Treatment with ruxolitinib was performed as previously described [2]. Briefly, Balb/c recipients were intravenously injected with $5x10^6$ allogeneic (C57BL/6J donor mice) T cell-depleted BM cells (TCD-BM) directly after myeloablative TBI with 2 x 4.5 Gy (medium dose). In some experiment, mice received 2 x 4.0 Gy (low dose) or 2 x 5.0-5.5 Gy (high dose) as indicated in the figure legends. Radiation was performed using the Gulmay RS225A irradiation device (Gulmay Medical, Camberley, UK) at a dose rate of 0.95 Gy/min (15 mA, 200 keV). Co-transplanation of allogeneic T cells was typically done with 0.5 x $10^6$ C57BL/6J purified donor T cells (medium dose). In some experiment mice received 0.1 (low dose), 1.5 - 2.5 x $10^6$ T cells respectively (high dose), as indicated in the figure legends. Donor T cells were isolated from pooled spleens of naïve C57BL/6J mice using a mixture of CD4 and CD8 MicroBeads (Order No: 130-117-043 and 130-117-044, Miltenyi Biotec). Bone marrow cells were isolated from naïve C57BL/6J mice and TCD was performed using CD90.2 MicroBeads (Order No: 130-121-278, Miltenyi Biotec). Individual experiments were performed with transplantations of BM and $T_{conv}$ cells (75-100 x $10^3$ $CD4^+$ $CD25^-$ + 75-100 x $10^3$ $CD8^+$ T cells) ± co-transfer of 150 x $10^3$ $T_{reg}$ cells ($CD4^+$ $CD25^+$). Cells were isolated using the Regulatory T Cell Isolation Kit (Miltenyi Biotec) according to the manufacturer's protocol and CD8 MicroBeads as described above. Weight loss was monitored at least once per week after allo-BMT. Mice received ruxolitinib (30 mg/kg body weight, purchased from Novartis under the brand name Jakavi®) dissolved in PEG300/dextrose 5% in a 1:3 ratio (PEG/dextrose) by oral gavage twice daily, starting from day -1 prior allo-BMT until the day before analysis.

##### Abdominal irradiation

Specific anatomic regions of mice were irradiated as previously described [3]. Mice were anesthetized with an intraperitoneal injection of medetomidin (0.5 mg/kg), midazolam (5 mg/kg), and fentanyl (0.05 mg/kg) and were fixed on their back on a plastic disc before irradiation. The entire abdominal area from the costal arch to the pelvis of the mice was then irradiated on 5 consecutive days (4.5 Gy per day, cumulative dose of 22.5 Gy) using the CIX2 irradiation device (Xstrahl). Lead plates (9 mm in total) were used to shield the rest of the body from radiation. Control mice were also anesthetized but did not receive irradiation.

**Irradiation of intestinal organoids and co-culture with $T_{reg}$ cells or cytokine stimulation**

Intestinal organoids were cultured and passaged as described above and irradiated using techniques as previously described [4]. Organoids were irradiated with 2 or 4 Gy dose of radiation using a Faxitron Cabinet X-ray System. Directly after irradiation, organoids were co-cultured with 100 x $10^3$ syngeneic CD25+ CD4+ Treg cells isolated as described above. Alternatively, organoids were stimulated with indicated cytokines after irradiation (0,25ng/mL recombinant murine IFNγ; 10ng/mL recombinant murine IL-10, both Peprotec). Cytokines or immune cells were removed after 4 days.

**Human organoid culture**

For human organoids, healthy tissue of colon resections of colorectal cancer patients was used. The tissue was cut in 5 mm pieces and incubated twice for 15 minutes in PBS + 30 mM EDTA. After washing with PBS, crypts were isolated by forcefully shaking for 30 seconds followed by 5-minute incubation on ice. This step was repeated four times. Afterwards crypts were strained through a 100 µm and a 80 µm strainer, embedded in Matrigel with 300 crypts per 50 µL drop and cultured in human organoid media [DMEM/F-12 (ThermoFisher), 10 mM HEPES, 2 mM L-Glutamin, 100 ng/mL human Noggin (Peprotec), 10 % human R-spondin conditioned media, 100 ng/mL murine Wnt3A (Peprotec), B27 supplement (Gibco), 1,25 mM N-Acetylcystein, 10mM Nicotinamid (Sigma), 50 ng/mL human EGF (Peprotec), 15 mM SB202190 (Sigma), 500 nM A83-01 (Sigma), antibiotic antimycotic solution (Sigma), 100 µg/mL Normocin (InvivoGen) and 10 µM Y-27632 (Sigma, only after seeding or passaging)].

The media was changed every 3 days and organoids passaged after 7 days. For experiments, established organoids were used from passages three and onwards.

**FITC Dextran in vivo permeability assay**

Experiments were performed as previously described [1]. Mice were kept without food and water for 6-8 h. Then, FITC-dextran (#FD4-1G, Sigma) was administered by oral gavage at a concentration of 75 mg/mL in water (750 mg/kg). 4 h later, plasma was collected from peripheral blood (8,800 rcf, 10 min), then mixed 1:1 with PBS and analyzed on a plate reader for FITC fluorescence signal at 490 nm excitation wavelength and 525 nm emission wavelength using the Varioskan™ LUX multimode microplate reader (Thermo Scientific).

**Leukocyte isolation from intestinal epithelium and lamina propria**

Isolation was performed similarly to previously described experiments [1,5]. Colon and/or ileum (defined as distal 1/3 of small intestine) were flushed with cold PBS and cut into 2 cm pieces. Longitudinally opened intestines were washed and then incubated with HBSS solution containing 2 mM EDTA, 10 mM HEPES, 5% FCS (Hyclone), 1% Penicillin-Streptomycin, 1% L-Glutamine and 1 mM DTT (all Sigma-Aldrich). After incubation on a shaker (200 rpm) at 37 °C for 15 min, tissues were washed and filtered through a 100 µm strainer (BD 352360), and flow through including the intraepithelial leukocytes (IEL) was placed on ice for 45 min. Next, intestines were incubated for 45 min in $PBS^{+Ca/+Mg}$ supplemented with FCS (10%), Collagenase II (200 U/mL; Worthington) on a shaker at 37 °C. Alternatively, digestion was performed with Liberase (400 U/mL) and DNAse1 (0.1 mg/mL) (Roche Diagnostics, Indianapolis, IN). Cells in suspension were filtered through a 100 µm strainer. LPL and IEL were purified on a 40/80% Percoll gradient (GE Healthcare Life Science, Pittsburgh, PA).

#### Flow cytometry

For intracellular cytokine staining, cells were stimulated for 3-4h with eBioscience™ Cell Stimulation Cocktail (plus protein transport inhibitors) (#00-4975-93). Single cell suspensions were stained with Live/Dead Yellow (Life Technologies, Grand Island, NY) and antibodies (Table S4) followed by fixation/permeabilization (Cytofix/Cytoperm, BD) and intracellular staining. Flow cytometry was performed using a Fortessa cytometer (BD) or CytoFLEX (Beckman Coulter) and data were analyzed with FlowJo 10 software (BD). *Ex vivo* analyses of very small cell populations (in particular large intestinal epithelial T cells) were performed as soon as at least 100 live $CD4^+$ cells and 10 live $CD4^+$ $Foxp3^+$ cells were identified.

#### Cell Sorting

For FACS purified $T_{reg}$ cell cultures, splenic $CD25^+$ cells from FoxP3-GFP reporter mice were enriched with anti-CD25-PE antibodies and anti-PE microbeads (Miltenyi Biotec), and were then stained for CD4. $T_{reg}$ cells were sorted on a FACS-ARIA II (BD Bioscience) as $CD4^+CD25^{high}FoxP3\text{-}GFP^+$. $T_{reg}$ cell purity was >98%.

Single cell suspensions of small intestinal organoids were generated by removing organoids from the culture with PBS and digestion with TrypleE Express for 10 minutes at 37°C. Single cells were passed through a 100µM strainer, washed and stained with EpCAM-BUV395 (BD Bioscience) and respective hashing antibody (TotalSeq-B0301-306 anti-mouse Hashtag 1-6 Antibody, Biolegend). Live/dead staining was performed immediately before sorting by adding propidium iodide. Single cells were sorted as $live/dead^-EpCAM^+$ cells.

#### Histological sample preparation

Mouse tissues were fixed in 10% (v/v) neutral-buffered formalin solution for a minimum of 48 h, dehydrated under standard conditions (Leica ASP300S, Wetzlar, Germany) and were embedded in paraffin. Serial 2 µm-thin sections prepared with a rotary microtome (HM355S, Thermo Fisher Scientific, Waltham, USA) were collected and subjected to histological and immunohistochemical analysis. Hematoxylin-Eosin (H.-E.) staining was performed on deparaffinized sections with Eosin and Mayer’s hemalum solution according to the standard protocol.

**Immunohistochemistry**

Immunohistochemistry was performed using a BondMax RXm system (Leica, Wetzlar, Germany, all reagents from Leica) with a primary antibody against CD3 (clone SP7, CI 597C01, DCS, Hamburg, Germany). In brief, slides were deparaffinized using deparaffinization solution, pretreated with Epitope retrieval solution 1 (corresponding to citrate buffer pH 6) for 30 minutes. Antibody binding was detected with a polymer refine detection kit without post-primary reagent and visualized with DAB as a dark brown precipitate. Counterstaining was done with hematoxyline.

**Histopathological evaluation**

Stained slides were scanned with an automated slide scanner (Leica Biosystems, Wetzlar, Germany, AT-2) and visually analyzed using the Aperio Imagescope software (version 12.3, Leica Biosystems, Wetzlar, Germany). The degree of $CD3^+$ T-cell infiltration was examined in a blinded fashion by an experienced pathologist. The degree of epithelial T cell infiltration was assessed by counting the number of infiltrating $CD3^+$ cells (intraepithelial lymphocytes) in 5 low power fields (20x magnification on scanned specimens) and calculating the average number of infiltrating cells for each specimen.

**RNA sequencing of bulk murine tissue**

<u>RNA isolation</u>: RNA was isolated from bulk tissue homogenates, which were prepared as follows: 1 cm of small intestine was flushed, and longitudinally slit pieces were frozen in 500 µL TRIzol (ambion) reagent using liquid nitrogen. After thawing, samples were homogenized using 5 mm stainless steel beads in a Tissue lyser II (Qiagen) for 1 min with 30 Hz (1800 oscillations/minute). Total RNA was isolated and used for RNA sequencing.

RNA sequencing: Library preparation for bulk 3'-sequencing of poly(A)-RNA was done as described previously[6]. Briefly, barcoded cDNA of each sample was generated with a Maxima RT polymerase (ThermoFisher) using oligo-dT primer containing barcodes, unique molecular identifiers (UMIs) and an adapter. 5' ends of the cDNAs were extended by a template switch oligo (TSO) and after pooling of all samples, full-length cDNA was amplified with primers binding to the TSO-site and the adapter. cDNA was tagmented with the Nextera XT kit (Illumina) and 3'-end-fragments finally amplified using primers with Illumina P5 and P7 overhangs. The library was sequenced on a NextSeq 500 (Illumina) with 16 cycles for the barcodes and UMIs in read1 and 65 cycles for the cDNA in read2.

Analysis of RNA sequencing: Gencode gene annotations version M18 and the mouse reference genome major release GRCm38 were derived from the Gencode homepage (https://www.gencodegenes.org/). Dropseq tools v1.12 [7] was used for mapping the raw sequencing data to the reference genome. The resulting UMI filtered count matrix was imported into R v3.4.4. Prefiltering the data was performed by calculating the median gene expression within each experimental condition. Genes having a groupwise median below 3 reads in at least 2 experimental conditions were filtered out. Prior differential expression analysis with Limma v3.40.6 [8], sample specific weights were estimated and used as coefficients alongside the T cell dosages as covariate during model fitting with Voom. The t-test was used for determining differentially regulated genes between all possible experimental groups. A gene was determined to be differentially regulated if the adjusted p-value was below 0.05. Gene set enrichment analysis was conducted with the preranked GSEA method [9] within the MSigDB Reactome, KEGG and Hallmark databases (v7.2). Genes were ranked according to their respective log2 fold change. A pathway was considered to be significantly associated with an experimental condition at an alpha level of 0.05. Genes that contribute most to the GSEA Enrichment Score for selected and significantly associated pathways for the comparison of the conditions 0.1 x $10^6$ T cells vs. 2.5 x $10^6$ T-Cell are visualized as Heatmap including all experimental conditions.

**Single cell RNA sequencing of human samples**

Single cell extraction from human gut biopsies: Biopsies were transferred into pre-warmed (37°C) HBSS (Hank's Balanced Salt Solution w/o $Mg^{2+}/Ca^{2+}$) as soon as possible after resection. Tissue pieces were disrupted mechanically, transferred into 5 mL of digestion buffer (25 mM HEPES, 0,1 % Collagenase IV, in HBSS w/o $Ca^{2+}/Mg^{2+}$), vortexed and digested in a thermo shaker (30 min, 37°C, 220 rpm, vortexed after 15 min). The resulting suspension was filtered using a 100 µm nylon-mesh into a 50 mL Falcon-tube, and the digestion was stopped by addition of 40 mL wash-buffer (RPMI1640, 10 % FCS, 25mM HEPES). Centrifugation (470x g, RT) was followed by another washing step with wash buffer, before the cells were resuspended in 250 µL freezing buffer (FCS, 10% DMSO) and frozen at -80°C (-1°C/min).

Thawing and sorting single cell suspensions from human gut biopsies: Each vial was thawed in 10 ml pre-warmed (37°C) RPMI and washed with 5 mL RPMI. Cell suspensions were stained on ice with CD45 (DAKO murine antiCD45 PB450, clone T29/33 # PB986) and Propidium iodide (PI). $CD45^{+}PI^{-}$ viable cells were sorted for scRNA sequencing.

10x experiments: After sorting, cells were centrifuged and the supernatant was carefully removed. Cells were resuspended in the Mastermix + 37.8 µL of water before 70 µL of cell suspension were transferred to the chip. (Step 1.1 and 1.2 of the original protocol). After each step, the integrity of the pellet was checked under the microscope to ensure that all cells are loaded onto the chip. From here on, 10x experiments have been performed according to the manufacturer's protocol (Chromium next GEM Single Cell VDJ V1.1, Rev D). QC has been performed with a High sensitivity DNA Kit (Agilent #5067-4626) on a Bioanalyzer 2100 as recommended in the protocol and libraries were quantified with the Qubit dsDNA hs assay kit (life technologies #Q32851). All steps have been performed using RPT filter tips (Starlab #S1183-1710, #SS1180-8710, #S1182-1730) and DNA LoBind tubes (Sigma #EP0030108051, #EP0030108078, #EP0030124359).

Sequencing: Libraries were pooled according to their minimal required read counts (20.000 reads/cell for gene expression libraries and 5.000 reads/cell for TCR libraries). Illumina paired end sequencing was performed with 150 cycles on a NovaSeq 6000.

Analysis of scRNA sequencing Data: Annotation was performed using cellranger (V3.0.2, 10x genomics) against the human reference genome GRCH38. All subsequent analysis has been performed using SCANPY[10]. After general preprocessing according to good practice in scRNA seq analysis (<10% mitochondrial genes, regressing out cell cycle, mitochondrial genes and total counts), data were count normalized per cell and logarithmized. $T_{reg}$ cells were identified via expression cut-offs of CD3 and FoxP3 genes.

**ChipCytometry**

ChipCytometry of human FFPE biopsies was performed as previously described [11]. Briefly, tissue sections were rehydrated on coverslips and antigen retrieval was performed using TRIS-EDTA buffer (pH 8.5). Sections were then transferred to CellSafe Chips (Zellkraftwerk) and IFNγ Fluorescent in-situ hybridization was performed following the multiplex V2 RNA Scope protocol (ACD Bio), and Opal 560 (Akoya bioscience) was used for the detection. The signal was inactivated using an oxidative quenching buffer (PBS, 24 mM NaOH, 4,5% $H_2O_2$), before cyclic immunofluorescence with photobleaching was performed on the chip.

**Single cell RNA sequencing of murine small intestinal organoids**

Murine small intestinal organoids were co-cultured with $CD25^{high}FoxP3\text{-}GFP^+$ $T_{reg}$ cells alone or in presence of IL-10 receptor and IFNγ receptor blocking antibodies, or were stimulated with IL-10 and IFNγ directly without $T_{reg}$ cell co-culture. Organoids were suspended to single cell suspensions and labeled with hash-tag oligo (HTO) antibodies to enable pooling (cell hashing) [12]. FACS was used to filter and sort a defined number of viable $EpCam^+$ intestinal epithelial cells as described above. All experimental groups were pooled in the process. Three experimental replicates with 7200 cells per treatment group were generated.

10x processing and sequencing

Each replicate was split into two batches of 18000 cells, and each batch was subjected to 10x Genomics processing to isolate cDNA of mRNA and HTOs of single cells using the Chromium Next GEM Single Cell 3' Reagent Kits v3.1 and 3' Feature Barcode Kit (Dual Index). The amplified libraries were then sequenced on an Illumina Nextseq 2000 device (P3 flow cell, all libraries multiplexed, 60% HTO libraries, 40% RNA-seq libraries) for initial probing. For one library, low average counts and a large degree of insufficient/ambiguous HTO labeling was observed, it was excluded from further processing. The remaining five libraries were subjected to deeper sequencing on an Illumina Novaseq 6000 device (S2 flow cell, all libraries multiplexed, 8% HTO libraries, 92% RNA-seq libraries). All sequencing runs were performed using the following parameters: PE-28-10-10-90. Reads from both sequencing runs were combined in the subsequent analysis.

Raw read processing

Base-calling and demultiplexing of sequencing reads was carried out using the bcl2convert provided by Illumina Dragon (v-3.8.4 NextSeq2000) and DragonServer (2.1 NextSeq 6000) (Illumina Inc., San Diego, California, USA) . Sequencing reads were combined and mapped to the mouse genome (https://cf.10xgenomics.com/supp/cell-exp/refdata-gex-mm10-2020-A.tar.gz", based on Mus_musculus.GRCm38.dna.primary_assembly.fa.modified and gencode.vM23.primary_assembly.annotation.gtf.filtered.) using cellranger (v 7.0.0) with standard options and expected cells set to 10000. Cellranger created count files for gene expression libraries and Cell-surface-marker (HTO) libraries.

Processing and analysis of single cell RNA-seq and HTO count data

Processing and analysis of single cell RNA-seq and HTO count data was mainly performed in R (version 4.1.1) with the *Seurat* package (version 4.0.5). Since cell hashing was applied, it was possible to pool different treatment groups in a single RNA-seq library. In order to assign treatment groups to single cells based on HTO counts, the *HTODemux()* function from the *Seurat* package was employed with default settings (positive-quantile parameter of 0.99) after centralized log normalization. Cell calling via *Cellranger* yielded a total of 45085 cells across the five libraries and all conditions. Cells that could not be assigned a treatment group due to insufficient HTO labeling (negative cells) were excluded from the analysis, resulting in 34507 remaining cells.

Doublets were identified based on ambiguous HTO labeling reported by *HTODemux()* and using the *scDblFinder* R-package (version 1.8.0, used together with *SingleCellExperiment*, version 1.16.0) and also excluded from the analysis[13]. *scDblFinder* was run in cluster-based mode using the top 30 principal components (PCs) associated with the 3000 top expressed genes to build the k-nearest-neighbor (kNN) network. The top 15 principal components were included when training the gradient boosting classifier. Three iterations of scoring were performed. Known doublets from cell hashing were supplied, but only used for score thresholding and not for training. To obtain clusters for use with *scDblFinder*, the *FindNeighbours()* and *FindClusters()* functions of *Seurat* were used with default parameters on each scRNA-seq library separately after excluding negative cells, using the top 30 principal components as input. The clustering resolution was 0.5. Clustering and data preparation for clustering are described in more detail in the following section. To obtain an estimate for the expected doublet rate to supply to *scDblFinder*, the probability $p_g$ that a doublet is formed from the same treatment group, as well as the probability $p_c$ that a doublet is formed from the same cluster (homotypic doublet) were estimated based on the respective group and cluster sizes. Then the expected doublet rate $r_e$ is estimated as $r_e = \frac{r_h \cdot (1 - p_c)}{1 - p_g}$, where $r_h$ is the observed doublet rate from cell hashing. However, *scDblFinder* is only very loosely bound to the $r_e$ values supplied. After removing doublets, 23496 cells remained.

In order to obtain a valid set of cells for analysis, cells have additionally been filtered to have a total RNA count between 10000 and 90000, a total number of features between 3000 and 10000 and a maximum percentage of mitochondrial genes of 8.5%. In this step, filtering removed about 35% to 60% of cells, depending on the library. One of the six libraries was excluded completely due to the very low number of UMI counts compared to the other libraries, and ambiguous cell hashing. Across all libraries, a total of 11301 cells were then used for subsequent analyses.

Counts were normalized for library size by dividing by the sum of total counts per cell and multiplying with a scale factor of $1 \cdot 10^6$. Afterwards, a value of 1 was added and the natural logarithm of the counts was computed. The 3000 top variable features were identified by applying *Seurat's FindVariableFeatures()* function with default parameters (variance stabilizing transformation as selection method). Data was then mean-centered, scaled to unit variance, and subjected to principal component analysis (PCA). The principal components (PCs) were then used as input to the *RunHarmony()* function from the *harmony* R-package (version 0.1.0) to integrate the data from the different libraries by removing batch effects. *RunHarmony()* was employed with default parameters, grouping by library.

Afterwards, *Seurat's FindNeighbours()* function was used with the first 30 dimensions of the *harmony*-transformed data as input to construct a nearest-neighbor graph. The *FindClusters()* function was then run with default parameters (standard Louvain algorithm) to perform graph-based clustering on the cells. Clustering was run at different resolutions to find the optimal clustering setup. For visualization, a UMAP (Uniform Manifold Approximation and Projection)[14] dimensionality reduction was additionally applied to the data, using *Seurat's RunUMAP()* function on the first 30 dimensions of the *harmony*-transformed data. A resolution of 1.0 was chosen for the final clustering setup, as it provided the best compromise between granularity and meaningfulness of clusters, as well as agreement with the localization of cells in UMAP space. There were no pronounced differences in clustering or UMAP projection across libraries.

For cell type annotation of single cells, the R-package *SingleR* (version 1.8.1) was used[15], together with scRNA-seq reference dataset from the murine intestine from Haber et al. (2017, GEO accession number GSE92332, full length atlas data)[16]. The reference labels taken from that data set were stem cells, TA (transient amplifying) cells, enterocyte progenitors (late and early were combined), enterocytes, enteroendocrine, goblet, paneth and tuft cells. As the method for determining reference label marker genes, the Wilcoxon rank-sum test was used, otherwise *SingleR* was run with default parameters. Briefly, *SingleR* computes correlations between cells from the query dataset and annotated cells from a reference dataset, based on a set of reference label marker genes inferred from the reference dataset. A given cell is then annotated with the reference label it correlates best with. To validate the *SingleR*-based annotation, scores of cell type marker gene signatures have additionally been computed for each cell, using the *UCell* R-package (version 2.0.1, together with R version 4.2.0)[17]. To this end, marker gene signatures for stem cells, enterocytes, goblet, paneth, enteroendocrine and tuft cells as well as cell cycle marker genes have been taken from Haber et al. (2017), where they are shown in Extended Data Figure 1. Additionally, cell cycle marker gene signatures specific to either G2M- or S-phase have been taken from *Seurat*, they are originally from Tirosh et al. (2016)[18]. *UCell* provides robust signature scores based on the Mann-Whitney U statistic[17]. It has been run with a maximum of 3000 ranked genes per cell, otherwise default parameters were employed. UMAP plots of single cells, colored by *UCell* scores of the given cell type signature, could then be inspected to derive possible cell type annotations of sets of cells, and be checked for agreement with the *SingleR*-based annotation.

When aligning *SingleR*-based annotation with clustering results in UMAP space, which can be seen in Figures S5A + B, respectively, enterocyte and enterocyte progenitor annotation for the large population in the center of the UMAP visualization, as well as goblet/paneth cell and enteroendocrine cell annotation is in acceptable agreement with the obtained clusters. Neither clustering nor annotation could distinguish goblet and paneth cells, though, and only a negligibly small number of cells resembled tuft cells according to annotation. Also, the clustering does not reflect the distinction between stem and TA cells, which can be seen as a trend from left to right (lower UMAP 1 to higher UMAP 1) in the large cell population in the center, and from top to bottom (higher UMAP 2 to lower UMAP 2) in the smaller population to the left – there, also enterocyte progenitors cannot be separated by the clustering. This distinction can also not be properly reproduced in clustering when increasing the resolution. However, when considering **Fig S5B**, marked clustering of the large center population in UMAP 2 direction, and of the smaller population on the left in UMAP 1 direction can be observed, which is not reflected by cell type annotation. Based on the UMAP plots of cell cycle signature *UCell* scores in **Fig. S6**, we assume that this clustering results from differences in cell cycle stage, where the upper-center population shows high G2M-phase signatures, while the lower-center population shows high S-phase signatures. Furthermore, stem, TA and enterocyte progenitor cells comprise both the large center population, as well as the smaller population to the left. We show in our study that the appearance of most cells in this population results from treatment.

Another reason why the clustering does not fully align with cell type annotation is that the transition from stem over TA to enterocyte progenitor cells would be expected to be smooth, with probably rather subtle differences. This is supported by the heatmap of centered and scaled marker gene expression in **Fig. S5E.** The columns correspond to single cells with *SingleR* celltype annotation, while the rows correspond to the expression of a given marker gene from a reference signature. While stem cells show a higher expression of stem cell markers than TA cells, their overall profile is relatively similar. A progression from stem cells over TA cells and enterocyte progenitors to enterocytes is perceivable. This transition is also visible in the heatmap of *SingleR* scores **(Fig. S5F).** The columns again correspond to single cells with *SingleR* celltype annotation, while the rows correspond to the *SingleR* scores for a given cell type label from the reference. Higher scores mean a higher correlation with and thus similarity to the reference label. *UCell* scores of marker gene signatures support the *SingleR* annotation. This can be seen in the UMAP plots of *UCell* scores **(Fig. S6)**: The stem cell score is highest in the rightmost part of the large center population as well as in the upper part of the smaller population to the left. It then diminishes from right to left and from top to bottom, respectively, indicating the aforementioned smooth transition. However, the smoothness of the transition also means that it is hard to define strict boundaries for stem, TA and enterocyte progenitor cells, and that the annotation is fuzzy/approximate. The enterocyte score is very high in the bottom population of the UMAP projection, matching the *SingleR* enterocyte annotation. It then diminishes upwards, indicating enterocyte progenitor cells. The leftmost part of the large center population as well as the bottom part of the smaller population to the left also show slightly higher enterocyte scores, which matches with the enterocyte progenitor annotation from *SingleR*. Of course, agreement between signature *UCell* scores and *SingleR* annotation is not unexpected, as the annotation of the scRNA-seq reference dataset from Haber et al. (2017) was done with these gene signatures. Nevertheless, provided the correctness of the reference dataset annotation, the match supports the validity of the *SingleR* annotation. The annotation obtained from *SingleR* was then used in subsequent analyses.

Differential gene expression and gene set enrichment analysis

For each population of cells annotated with a given cell type (except tuft cells, due to the very low number of cells), testing for differential gene expression between treatment groups was performed on the raw count data with the *NEBULA* R-package (version 1.2.2)[19]. *NEBULA* fits (by default) a negative binomial mixed model to the count data, and estimates both cell- and subject-level overdispersions. For each cell type, a *NEBULA* negative binomial gamma mixed model was fitted with treatment group and library (batch) as categorical variables, as well as the number of features per cell as a continuous variable (to account for possible effects not captured by a library size scaling factor). Each combination of treatment group and experimental replicate was considered a distinct subject. Library size was set as the scaling factor, and minimum counts per cell for a gene to be tested were set to 0.005. The fit was computed with the *NEBULA*-LN method. All other parameters were set to default values. In order to be able to make all desired comparisons, contrasts were employed. To that end, the covariance matrix of the model was extracted and used together with the log fold-change (log FC) estimates and a vector of linear contrasts in a chi-squared test, as described in the vignette of the *NEBULA* package.

The differential gene expression results for a given cell type and comparison of treatment groups were then used as input to pathway/gene set analysis, using the GSEA (gene set enrichment analysis) approach as implemented in the *fgsea* R-package (version 1.20.0)[9]. As input for GSEA, the negative $\log_{10}$ p-values signed with the direction of the log FC were used. If there were p-values with a value of zero, these were set to the smallest non-zero normalized floating-point number ($2.23 \cdot 10^{-308}$) prior to log-transformation. Values were then sorted by the signed log-transformed p-values in descending order, breaking possible ties by sorting according to log FC. Gene sets were obtained from MSigDB (Molecular signatures database) [20] via the *msigdbr* R-package (version 7.4.1). Chosen gene set collections were hallmark gene sets (H), the KEGG and Reactome subsets from curated gene sets (C2), as well as transcription factor targets from regulatory target gene sets (C3). Only gene sets where at least 70% of genes were in the genes tested for differential expression in the given cell type were assessed. For each cell type, the p-values obtained from GSEA for the different comparisons and gene sets were corrected for multiple testing with the false discovery rate (FDR) approach[21], controlling for an FDR of 10%. In order to assess the similarity of treatment conditions in terms of pathway activation, the GSEA-derived normalized enrichment scores (NES) of different comparisons (within each cell type) have been correlated using the Pearson correlation coefficient. To reduce statistical noise, only gene sets significantly regulated at an FDR of 10% in either of the two comparisons have been used in the correlation.

Analysis of IFNγ and IL-10 gene expression in tissue $T_{reg}$ cells

For analyzing gene expression of IFNγ and IL-10 in tissue $T_{reg}$ cells, a published a scRNA-seq dataset (GEO accession number: GSE223798) was used. Briefly, donor $T_{reg}$ cells were expanded *in vitro*, supplied to recipient mice and then extracted from the respective tissues to assess tissue adaption [22]. Testing for differential gene expression between $T_{reg}$ cell populations from different tissues (or input $T_{reg}$ cells) was performed with the Wilcox test, via the *FindMarkers()* function of the *Seurat* package (version 4.0.5). Resulting p-values were corrected for multiple testing with the Bonferroni method.

Additional software/packages

Other R-packages used were ggplot2 (version 3.3.5), patchwork (version 1.1.1), dplyr (version 1.0.7) stringr (version 1.4.0) and tidyr (version 1.1.4), pheatmap (version 1.0.12) and viridis (version 0.6.2), as well as foreach (version 1.5.1), doParallel (version 1.0.16), future (version 1.31.0), rprojroot (version 2.0.2), yaml (version 2.2.1) and WriteXLS (version 6.3.0). For creating Venn diagrams, Python (version 3.9.17) was used together with the packages numpy (version 1.25.2), pandas (version 2.0.3), matplotlib (3.7.1) and matplotlib-venn (0.11.9).

**II. Table S4**

| Antibodies | | |
|---|---|---|
| REAGENT | SOURCE | IDENTIFIER |
| Anti-human Pan-Cytokeratin (AF488) | BioLegend | RRID: AB_2616664 |
| Anti-human CD4 (AF488) | R&D systems | RRID: AB_2728839 |
| Anti-human Foxp3 (PE) | Thermo Fisher Scientific | RRID: AB_1944444 |
| Anti-human CD45 (PerCP/Cy5.5) | BioLegend | RRID: AB_893338 |
| Anti-human CD3 | Thermo Fisher Scientific | RRID: AB_149924 |
| Anti-mouse CD3-AF488 | Biolegend | RRID: AB_389301 |
| Anti-mouse PD-1 (PE) | Biolegend | RRID: AB_1877231 |
| Anti-mouse CD4 (PerCp/Cy5.5) | Biolegend | RRID: AB_893324 |
| Anti-mouse CD8 (BUV395) | BD | RRID: AB_2732919 |
| Anti-mouse H2K-B (BUV421) | Biolegend | RRID: AB_2876430 |
| Anti-mouse CD45 (AF700) | Biolegend | RRID: AB_493715 |
| Anti-mouse IFNγ (PE-Cy7) | Biolegend | RRID: AB_2295770 |
| Anti-mouse Foxp3 (AF647) | Biolegend | RRID: AB_439750 |
| Anti-rabbit IgG (PE) | Biolegend | RRID: AB_2563484 |
| Anti-mouse CD45 (APC-Cy7) | Biolegend | RRID: AB_312981 |
| Anti-mouse Foxp3 (PE) | Biolegend | RRID: AB_1089117 |
| Anti-mouse CD16/32 Fc block | Biolegend | RRID: AB_2783138 |
| Anti-mouse IL-10 (BV421) | Biolegend | RRID: AB_2563240 |
| Anti-Mouse CD326 (BUV395) | BD | RRID: AB_2740020 |
| TotalSeq™-B0301 anti-mouse Hashtag 1 Antibody | Biolegend | RRID: AB_2814067 |
| TotalSeq™-B0302 anti-mouse Hashtag 2 Antibody | Biolegend | RRID: AB_2814068 |
| TotalSeq™-B0303 anti-mouse Hashtag 3 Antibody | Biolegend | RRID: AB_2814069 |
| TotalSeq™-B0304 anti-mouse Hashtag 4 Antibody | Biolegend | RRID: AB_2814070 |
| TotalSeq™-B0305 anti-mouse Hashtag 5 Antibody | Biolegend | RRID: AB_2814071 |
| TotalSeq™-B0306 anti-mouse Hashtag 6 Antibody | Biolegend | RRID: AB_2814072 |

### III. Supplemental Figures

#### Supplemental Figure 1

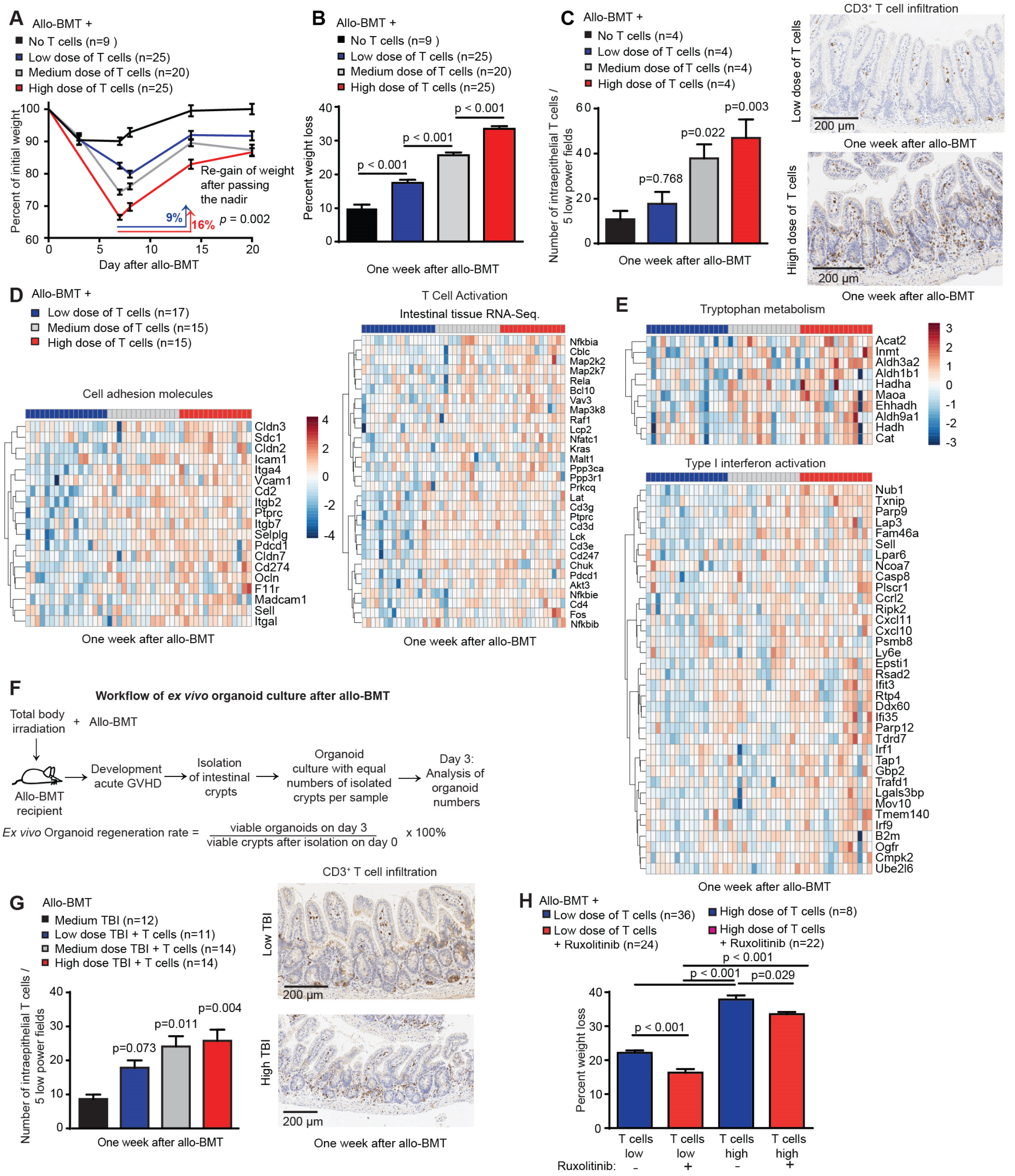


**Intestinal T cell infiltration determines acute weight loss and regeneration after allo-BMT**

**A)** Weight loss of Balb/c mice that received 9 Gy TBI followed by allo-BMT (C57BL/6J donors) of BM ± allogeneic T cells (low dose: 0.1 x10$^6$ T cells; medium dose: 0.5 x10$^6$ T cells; high dose: 2.5 x10$^6$ T cells). The depicted regain of weight was calculated between initial weight loss on day 7 and recovery until day 14 after allo-BMT. Pooled data of 3 independent experiments. **B)** Maximal weight loss one week after allo-BMT. Pooled data of three independent experiments. **C)** Immunohistochemical analysis of $CD3^+$ intestinal intraepithelial T cells one week after allo-BMT with 9 Gy TBI. Representative images of intraepithelial T cell infiltration with low dose or high dose of T cells. **D-E)** Next generation RNA sequencing from bulk small intestinal tissue one week after allo-BMT: gene set enrichment analysis was conducted with the preranked GSEA method within the MSigDB Reactome, KEGG and Hallmark databases. Genes that contribute most to the GSEA Enrichment Score for the selected pathways (Cell Adhesion Molecules and tryptophan Metabolism) of the KEGG database for the comparison of low vs. high T cells doses are shown as heatmap including all three experimental T cell conditions. **F)** Workflow of *ex vivo* organoid culture: on day 7 after allo-BMT, recipient mice were sacrificed, small intestinal crypts isolated and in constant numbers were used for intestinal organoid culture. Organoids were counted on day 3 of culture. **G)** Analysis of $CD3^+$ intestinal intraepithelial T cells one week after allo-BMT. Low (8Gy) vs. high (10 Gy) dose TBI + BM + T cells were compared using the one-tailed unpaired t-test. Representative images of T cell infiltration. **H)** Balb/c mice received TBI (9 Gy) followed by allo-BMT with BM and a low ($0.1 \times 10^6$) or high dose ($2.5 \times 10^6$) of T cells +/- ruxolitinib treatment. Peak weight loss of recipients one week after allo-BMT. Data are presented as mean ± S.E.M. Data were analyzed using ordinary one-way ANOVA for multiple comparisons unless otherwise stated above. Animal numbers per group (n) are depicted.

**Supplemental Figure 2**

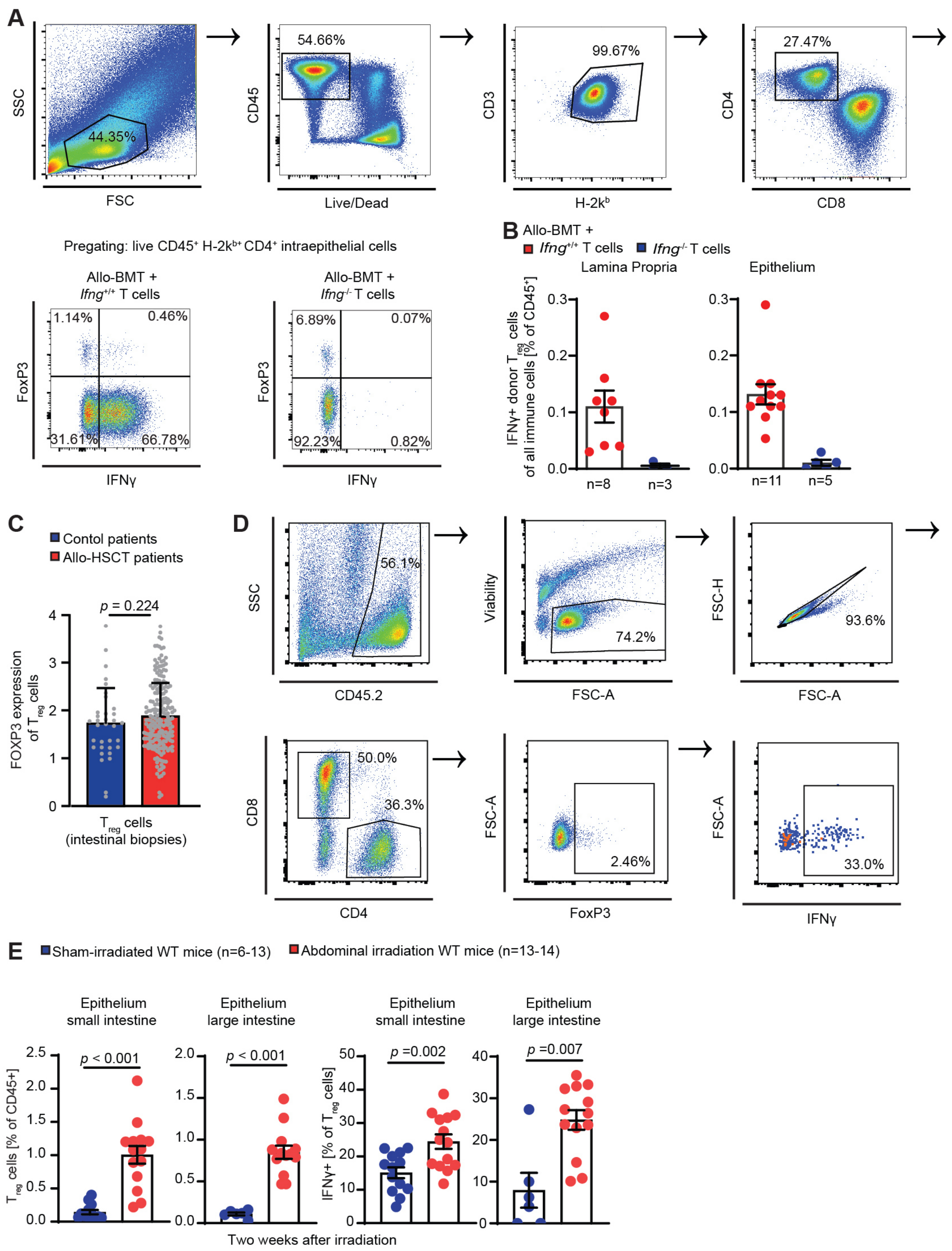


**Enhanced intestinal abundance of IFNγ-expressing intestinal $T_{reg}$ after tissue injury**

**A)** Balb/c mice received TBI (9 Gy) followed by allo-BMT with BM and a high dose of T cells derived from either WT or IFNγ-deficient (IFNγ$^{-/-}$) donor animals (C57BL/6). Gating strategy and representative FACS Gating of IFNγ$^{+}$ FoxP3$^{+}$ CD4$^{+}$ donor (H-2kb$^{+}$) cells within all CD45$^{+}$ intestinal intraepithelial or lamina propria leukocytes. **B)** Frequency of IFNγ$^{+}$ FoxP3$^{+}$ CD4$^{+}$

donor (H-2kb$^+$) cells within all CD45$^+$ intestinal intraepithelial or lamina propria leukocytes analyzed by flow cytometry. Pooled data of three independent experiments. **C)** *FOXP3 gene* expression of $T_{reg}$ cells were analyzed via scRNA-Seq of cells isolated from large intestinal biopsies of allo-HCST recipients (n=22 patients, with n=208 identified $T_{reg}$ cells) or control patients that did not undergo allo-HSCT (n=5 patients with n=34 identified $T_{reg}$ cells). **D)** C57BL/6 WT mice received abdominal irradiation (ABI, 5 x 4,5 Gy/day from day 0 until day 4). Gating strategy for flow cytometry of intraepithelial leukocytes of the small intestine two weeks (d15) after start of ABI in a WT mouse. **E)** Data shown the percentage of $T_{reg}$ cell of all live immune cells and the percentage of IFNγ$^+$ of all $T_{reg}$ cells in mice ± ABI. Pooled data from 3 independent experiments. All data were analyzed using unpaired t-test, Mann–Whitney U test (Fig. S2C) or ordinary one-way ANOVA for multiple comparisons and are presented as mean ± S.E.M. Animal numbers per group (n) are depicted.

#### Supplemental Figure 3

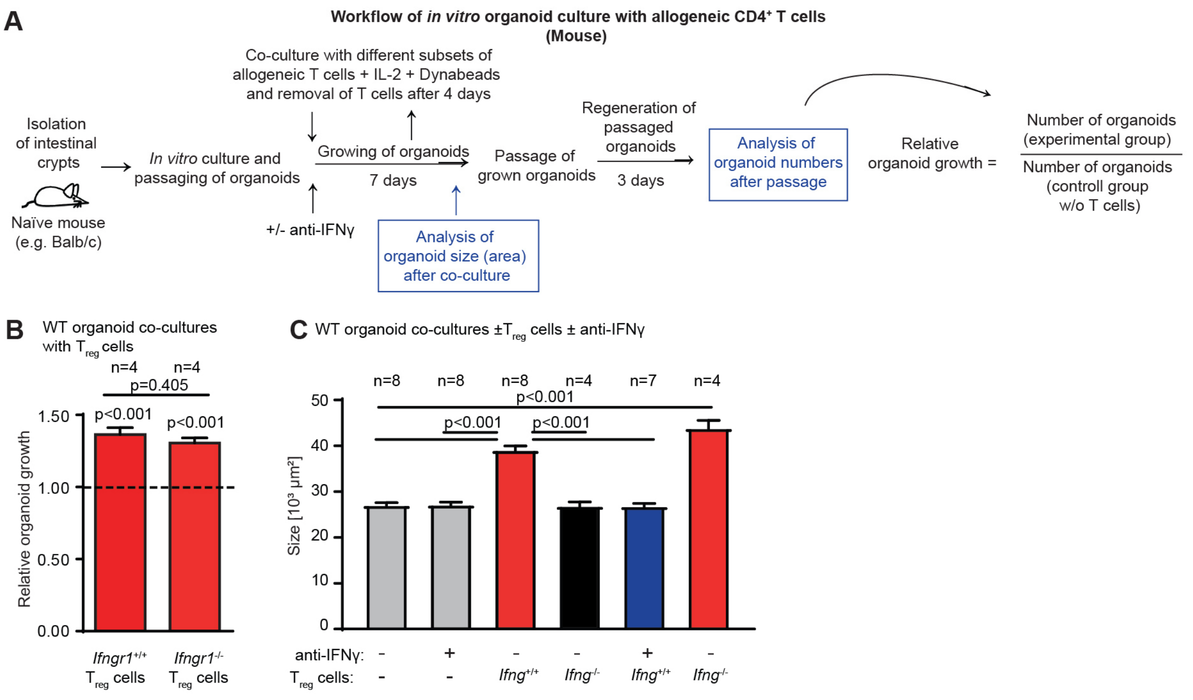


**$T_{reg}$ cell-derived IFNγ promotes the growth of intestinal organoids independently of an IFNγR feedback loop on $T_{reg}$ cells**

**A)** Work flow of *in vitro* organoid coculture as described in Material and Methods. **B)** Relative organoid growth of small intestinal organoids co-cultured with allogeneic $T_{Reg}$ cells isolated from WT (IFNγR$^{+/+}$) or IFNγ receptor-deficient (IFNγR$^{-/-}$) mice. **C)** Mean size (area) of organoids on day 6 after coculture with allogeneic $T_{Reg}$ cells. Cultures were stimulated as described above (IL-2, +/- anti-IFNγ) and $T_{Reg}$ cells were isolated from indicated donor animals (wild-type, IFNγ$^{-/-}$, IFNγR$^{-/-}$). Pooled data from all performed experiments described above with the indicated experimental groups (number of measured organoids: untreated (n=1129); anti-IFNγ (n=772); WT $T_{Reg}$ (n=656); IFNγ$^{-/-}$ $T_{Reg}$ (n=349); WT $T_{Reg}$ + anti-IFNγ (n=579); IFNγR$^{-/-}$ $T_{Reg}$ (n=280). The number (n) of separate organoid culture experiments is indicated in the figure.

**Supplemental Figure 4**

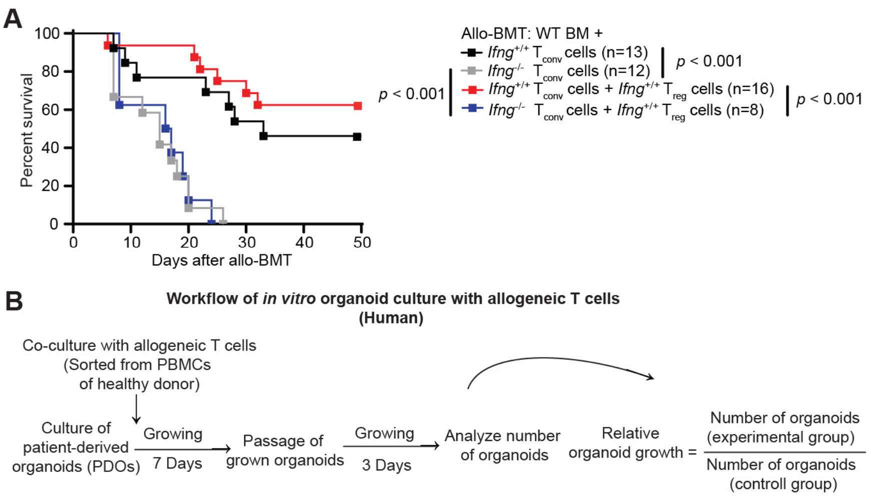


**$T_{conv}$ cell derived IFNγ is essential for $T_{reg}$ cell mediated protection from GVHD after allo-BMT.**

**A)** Balb/c mice received TBI (9 Gy) followed by allo-BMT with WT BM and IFNγ$^{+/+}$ or IFNγ$^{-/-}$ $T_{Conv}$ cells (C57BL/6J donors). Indicated mice received a co-transfer of IFNγ$^{+/+}$ $T_{Reg}$ cells. Pooled data from two independent experiments. **B)** Workflow of *in vitro* human organoid coculture with allogeneic T cells: Large intestinal organoids (patient-derived organoids, PDOs) were co-cultured with allogeneic T cells. Three days after a passage, the number of established organoids were counted and analyzed. Relative organoid growth is normalized to the number of organoids in steady-state culture without T cells. Animal numbers per group (n) are depicted.

#### Supplemental Figure 5

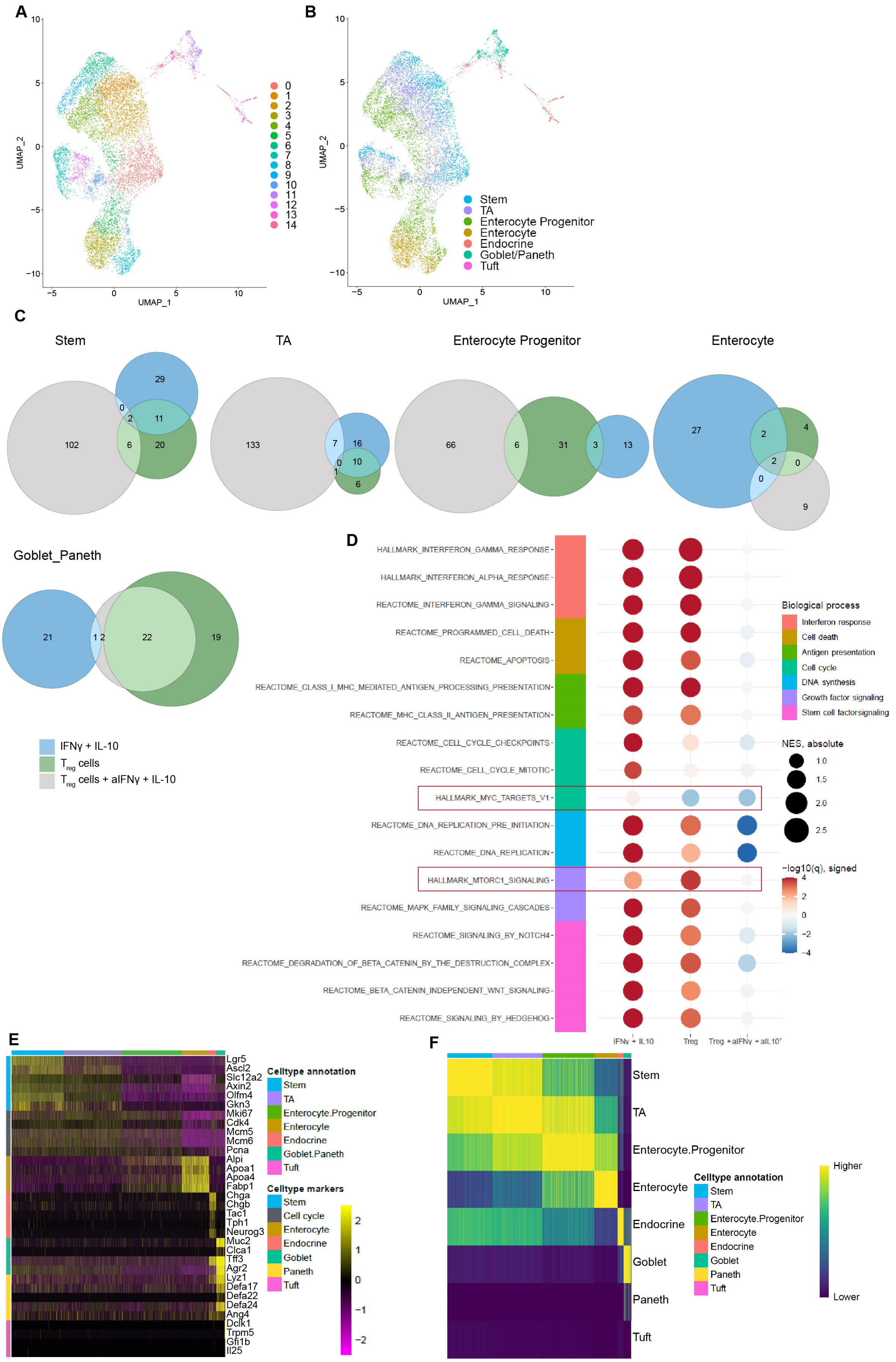

**Clustering, cell type annotation and overlap of downregulated signaling pathways between experimental conditions**

**A)** Single cell RNA sequencing of murine small intestinal organoids after $T_{Reg}$ cell coculture or cytokine stimulation as described for Figure 5. UMAP plot of clustering results with a resolution of 1.0. Plot of single cells in UMAP space, colored by the results of graph-based clustering. **B)** UMAP plot of SingleR cell type annotation. Plot of single cells in UMAP space, colored by SingleR cell type annotation. **C)** Venn diagrams indicating the overlap of downregulated pathways between experimental conditions compared to control organoids. Only gene sets/pathways significantly upregulated when controlling for an FDR of 10% were considered. **D)** Dotmap of GSEA results of selected pathways/gene sets for different treatments (vs. control), enterocyte progenitor cells. Dots are colored by the negative $\log_{10}$ of the GSEA q-value (FDR), the sign indicates the direction of the regulation (up positive, down negative). The size of the dots corresponds to the GSEA NES. Gene sets/pathways are derived from the Hallmark (H) and Reactome (R) gene set collections of MSigDB. **E)** Heatmap of marker gene expression. Columns correspond to single cells, the bar on top provides the cell label obtained from SingleR annotation. Rows correspond to the gene expression of marker genes, where the bar to the left groups the marker genes according to their respective cell type/process. Gene expression is shown mean-centered, scaled to unit variance and clipped at values -2.5 and 2.5. **F)** Heatmap of SingleR scores. Columns correspond to single cells, the bar on top provides the cell label obtained from SingleR annotation. Rows correspond to the SingleR scores for a given cell type label from the reference. Higher scores denote a higher correlation with and thus similarity to the reference label. Scores of single cells have been min-max normalized to lie within a [0, 1] interval, and transformed to the power of 3 to improve visibility of the dynamic range near 1 (see documentation of the SingleR package).

#### Supplemental Figure 6

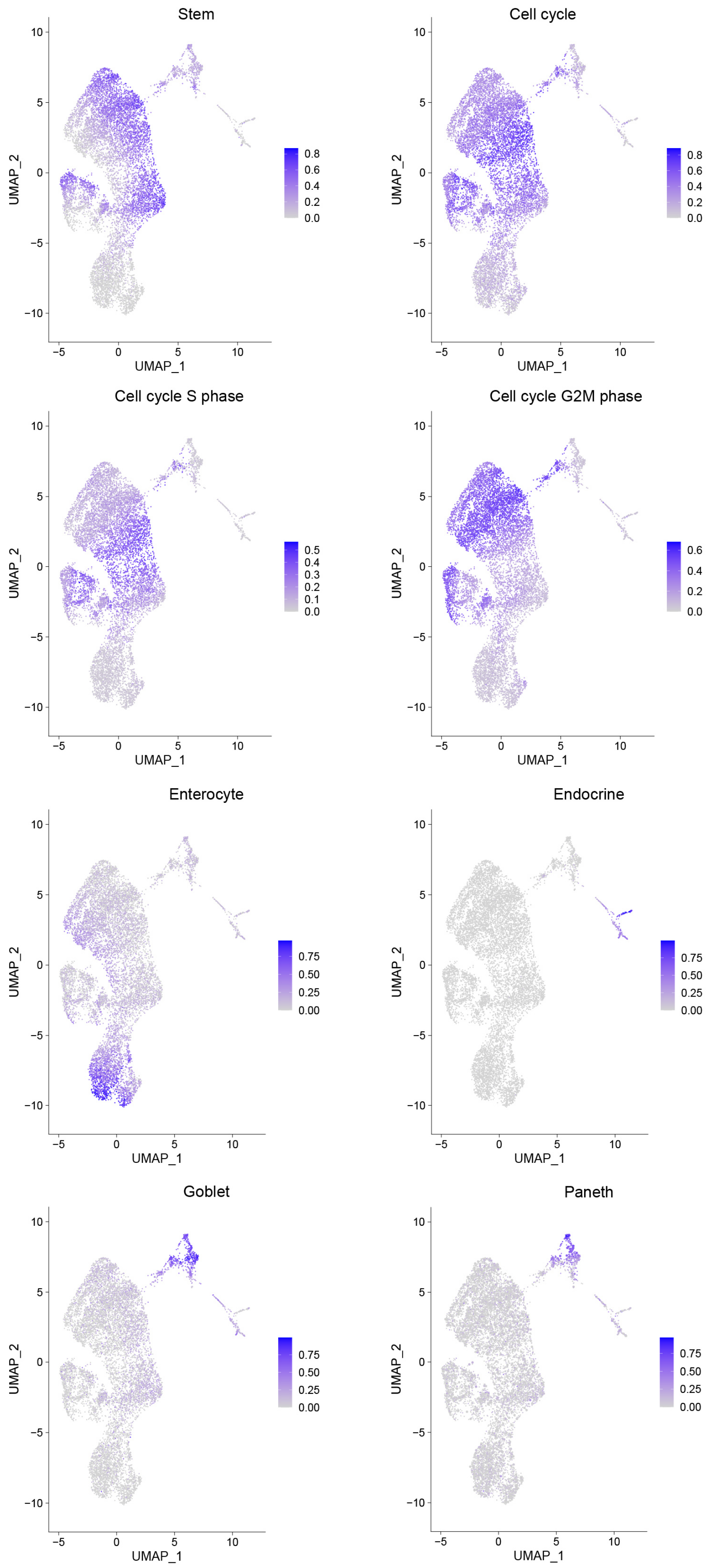

**UMAP plots of UCell scores and pots of single cells in UMAP space of single cell RNA sequencing analysis of murine small intestinal organoids**

UMAP plots of UCell scores. Plots of single cells in UMAP space, colored by the UCell scores of the respective marker gene signature.
